# *APOK3*, a pollen killer antidote in *Arabidopsis thaliana*

**DOI:** 10.1101/2021.09.08.459474

**Authors:** Matthieu Simon, Stéphanie Durand, Anthony Ricou, Nathalie Vrielynck, Baptiste Mayjonade, Jérôme Gouzy, Roxane Boyer, Fabrice Roux, Christine Camilleri, Françoise Budar

## Abstract

According to the principles of heredity, each parental allele of hybrids equally participates in the progeny. At some loci, however, it happens that one allele is favored to the expense of the other. Gamete killers are genetic systems where one allele (the killer) triggers the death of the gametes carrying the other (killed) allele. They have been found in many organisms, and are of major interest to understand mechanisms of evolution and speciation. Gamete killers are particularly prevalent in plants, where they can compromise crop breeding. Here, we deciphered a pollen killer in *Arabidopsis thaliana* by exploiting natural variation, *de novo* genomic sequencing and mutants, and analyzing segregations in crosses. We found that the killer allele carries an antidote gene flanked by two elements mandatory for the killing activity. We identified the gene encoding the antidote, a chimeric protein addressed to mitochondria. This gene appeared in the species by association of domains recruited from other genes, and it recently underwent duplications within a highly variable locus, particularly in the killer genotypes. Exploring the species diversity, we identified sequence polymorphisms correlated with the antidote activity.

## Introduction

Genetic loci that do not comply with Mendel’s laws have been observed since the dawn of genetics. First considered as being genetic curiosities, these loci with transmission ratio distortion (TRD) are now recognized to be common in fungi, plants and animals, with a particularly high incidence in plants (Fishman and McIntosh 2019). They are of major interest to understand genomic evolution, adaptation and speciation (Presgraves, 2010; Lindholm et al, 2016; Fishman & Sweigart, 2018; Agren & Clarck, 2018). In addition, they have significant impacts on plant breeding, disturbing QTL mapping and hampering the use of genetic resources. Cases of TRD have been reported in multiple plant species, with most studies conducted in rice and in *Arabidopsis thaliana*. In rice, TRD is very common in inter-specific and sub-specific crosses, and often linked to hybrid sterility (Ouyang and Zhang 2018), limiting intersubspecific crosses suitable for breeding (Matsubara et al. 2011; Zhang et al. 2020). For instance, 18 genomic regions were recently found to be distorted in a cross between *Oryza sativa* ssp *japonica* and *O. sativa* ssp *indica* (Zhang et al. 2020). In the genus Arabidopsis, many examples of inter or intra-specific hybrid incompatibilities were also reported (reviewed in Vaid and Laitinen, 2019). In *A. thaliana*, TRD was observed in over half of a set of 17 F2 populations (Salomé et al, (2012). More recently, at least one TRD was detected in ∼25% of a set of 500 F2 populations, corresponding to more than one hundred distorted genomic regions (Seymour et al. 2019).

Causes of TRD are often classified into two main non-exclusive types, *i*.*e*. (i) Bateson-Dobzhansky-Muller (BDM) incompatibilities, where independently evolved alleles result in deleterious or sub-optimal phenotypes when brought together, and (ii) allele specific gamete elimination, where one allele takes over the alternative allele in the gametes produced by a heterozygote (reviewed in Ouyang and Zhang, 2013). Allele-specific gamete elimination can occur at the meiotic stage, hence designated meiotic driver. It is the case, for example, of the B chromosomes of cereals (Östergren 1945; Houben 2017) and the driving centromere in yellow monkeyflower (Finseth et al. 2015; Finseth et al. 2021). Alternatively, gamete elimination occurs after meiosis when the favoured (killer) allele induces a defect in the gametes that carry the alternative (killed) one, eventually causing their underrepresentation in the next generation. Such a situation was observed for the *wtf* genes in fission yeast (Nuckolls et al. 2017), the *Spok* genes in *Podospora anserina* (Grognet et al. 2014), the *SD* system in *Drosophila melanogaster* (Larracuente and Presgraves 2012), and the Sa (Long et al. 2008) and S5 (Yang et al. 2012) loci in rice. In plants, loci causing TRD by allele specific gamete elimination are designated gamete killers, or more specifically pollen killers (PK) when they affect the male gametes. In the unraveled TRD cases in *A. thaliana*, the identification of causal genes showed that the distortions resulted from BDM incompatibilities between parental alleles (or epialleles), most often located at physically unlinked loci (Bomblies et al. 2007; Bikard et al. 2009; Durand et al. 2012; Agorio et al. 2017; Jiao et al. 2021), or at one locus (Smith et al, 2011). However, in contrast to rice where several gamete killers have been studied (Ouyang and Zhang 2013), PKs have been reported only once in *A. thaliana* (Simon et al, 2016), and to our knowledge none has been molecularly deciphered so far in this species.

Meiotic drivers and gamete killers have retained particular attention for their potential in triggering genomic conflicts (Lindholm et al. 2016; Agren and Clark 2018). The most intriguing feature of gamete killers is that killer alleles trigger a defect in the gametes that do not carry them. This can be explained by one of two main genetic models (Bravo Nunez et al. 2018), schematically represented in Figure 1. In both models, all the genes responsible for gamete elimination are tightly linked and the killing factor is produced before (or during) meiosis, and still present in all the developing gametes. In the ‘killer-target’ model, a partner (the target), necessary for the killing activity, is encoded by the killed allele and expressed after meiosis, thus present only in gametes that carry it. This model applies to the *SD* system of *D. melanogaster* and the Sa locus of rice, for example (Bravo Nunez et al. 2018). In the ‘poison-antidote’ model, the killer allele also produces an antidote that counteracts the poison, but whereas the poison produced before meiosis subsists in the gametes, the antidote does not, so only the gametes that are able to produce it are rescued. Poison-antidote type segregation distorters have been described in fungi and plants and include the fission yeast *wtf* genes (Nuckolls et al. 2017) and the rice *qHMS7* PK (Yu et al. 2018).

**Figure 1:**
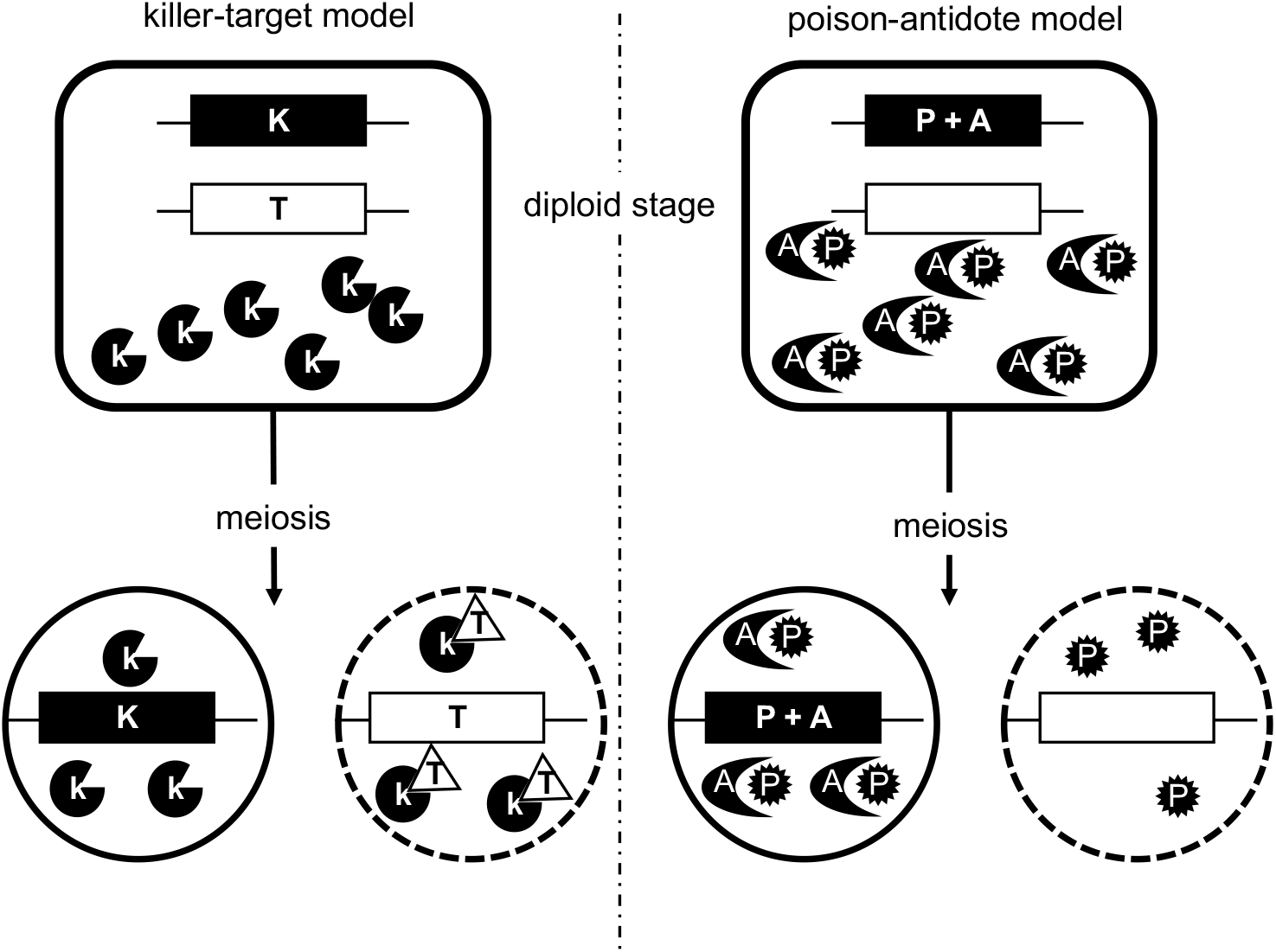
Two genetic models of gamete killers. The locus is represented as a horizontal line with the killer and killed alleles as black and white boxes, respectively. For these general models, there is no hypothesis on the number of genes present in each box. In both models, the black allele is expressed before meiosis, and the killer (K) or poison (P) persists in all the gametes. In the killer-target model, only gametes with the white allele express the target (T), which interacts with K to trigger gamete death (represented as dashed outline). In the poison-antidote model, the black allele also produces a short-lived antidote (A) that counteracts P, so only the gametes that inherit the black allele are efficiently protected.

Here we decipher one of the PKs that were uncovered to contribute to a male sterility observed in hybrids between two *A. thaliana* natural variants, Shahdara (Sha) and Mr-0 (Simon et al, 2016). We show that this PK, located at the bottom of chromosome 3, belongs to the poison-antidote class, and we identified the antidote gene. This gene is chimeric and encodes a mitochondrial protein; it originated and evolved in the species *A. thaliana* within a highly variable locus. We precisely characterized the locus, showing that it contains at least two killer elements required for its activity, flanking the antidote gene. By exploring the diversity within the species, we found this PK in a number of *A. thaliana* hybrids. We then showed that in the killer genotypes, the locus strongly differs from those of neutral and killed genotypes by important structural variations, including duplications of the antidote gene. Lastly, we identified in the antidote several polymorphisms that correlate with its protective activity.

## Results

### Elements leading to a segregation bias at the L3 locus are common in natural variants of *A. thaliana*

Amongst the PKs detected in the selfed progeny of Mr-0 x Sha hybrids (Simon et al. 2016), the L3 locus, located at the bottom of chromosome 3, induced a deficit in Sha homozygous progenies. This was linked to the death of pollen grains carrying the Sha allele at this locus. We also observed a strong bias at L3 against the Rak-2 allele in the Mr-0 x Rak-2 F2 population (Simon et al. 2016), suggesting that some natural variants other than Sha possess at L3 alleles sensitive to the Mr-0 killer effect. Likewise, other variants than Mr-0 could carry killer alleles. After crossing 30 accessions belonging to different diversity groups (Simon et al. 2012) to Mr-0 and/or Sha, we analyzed the segregation at L3 in the progeny of the hybrids to reveal a possible killed or killer behaviour. Similar to Sha, 14 of the 26 accessions tested in crosses with Mr-0 for the killed status showed a bias in the progeny of their hybrid (Table 1). Genotype proportions were consistent with a 1:1 distribution of homozygotes Mr-0 and heterozygotes in most of the biased segregations, as expected if there is a gametophytic defect in these hybrids (Table1 Source Data 1). On the other hand, among 18 accessions analyzed for a killer behaviour, five induced a bias in the progeny of their hybrid with Sha, as did Mr-0 (Table 1), with a distribution of genotypes that was consistent with a gametophytic defect (Table1 Source Data 2). Among the 14 accessions that were tested for both killed and killer status, five accessions, including Col-0, had a neutral behaviour (neither killed nor killer). And, coherently, none of these 14 accessions was found to have both a killed and a killer behaviour (Table 1).

**Table 1:** Killed or killer behaviors of natural accessions

| Accession | Country | Bias in cross with<br>Mr-0: killed<br>behaviour<sup>(1)</sup> | Bias in cross with<br>Sha: killer<br>behaviour<sup>(2)</sup> |
| --- | --- | --- | --- |
| **Shigu-2** | Russia | No | Yes |
| **Cant-1** | Spain | No | Yes |
| **Etna-2** | Italy | No | Yes |
| **Ct-1** | Italy | No | Yes |
| **Jea** | France | No | Yes |
| **Lov-5** | Sweden | No | No |
| **Blh-1** | Czech Republic | No | No |
| **Col-0** | Poland | No | No |
| **Bur-0** | Ireland | No | No |
| **Oy-0** | Norway | No | No |
| **Koch-1** | Ukraine | No | nd |
| **N16** | Russia | No | nd |
| **Sorbo** | Tajikistan | Yes | No |
| **Cvi-0** | Cape Verde Islands | Yes | No |
| **Ita-0<sup>(3)</sup>** | Morocco | Yes | No |
| **Are-10** | Portugal | Yes | No |
| **Are-1** | Portugal | Yes | nd |
| **Kas-2** | India | Yes | nd |
| **Kz-1** | Kazakhstan | Yes | nd |
| **Kidr-1** | Russia | Yes | nd |
| **Kly-1** | Russia | Yes | nd |
| **N13** | Russia | Yes | nd |
| **Nov-01** | Russia | Yes | nd |
| **Rak-2** | Russia | Yes | nd |
| **Stepn-1** | Russia | Yes | nd |
| **Zal-3** | Kyrgyzstan | Yes | nd |
| **Kz-9** | Kazakhstan | nd | No |
| **Kas-1** | India | nd | No |
| **Qar-8a** | Lebanon | nd | No |
| **Etn-0** | Italy | nd | No |

<sup>(1)</sup> Table1-Source Data 1: Segregations at L3 in progenies of crosses of Mr-0 with different natural accessions. <sup>(2)</sup> Table1-Source Data 2: Segregations at L3 in progenies of crosses of different natural accessions with Sha. <sup>(3)</sup> Ita-0 was tested by crossing not with Mr-0 but with two other killer accessions, Ct-1 and Jea. nd: not determined.

Therefore, this segregation distorsion at L3 appears to be widespread in crosses between natural accessions of *A. thaliana*, and we focused on the Mr-0 x Sha cross to investigate the underlying genetic elements.

### The segregation distorter at L3 in Mr-0 x Sha hybrids induces an allele specific impaired pollen development

We previously showed that a plant segregating Sha and Mr-0 alleles only at L3 while fixed Sha in the rest of its nuclear genome (hereafter ShaL3^H^) presented a strong bias against Sha homozygous progenies, which was linked to a deficit in pollen grains carrying the Sha allele (Simon et al. 2016). Here, we tested whether the segregation bias was dependent on the genetic background by comparing the selfed progenies of ShaL3^H^ with its equivalent in the Mr-0 nuclear background (MrL3^H^). Their progenies showed very similar deficits in Sha homozygotes (Table 2), indicating that the segregation distortion was independent of the fixed parental nuclear background. Accordingly, anthers from both genotypes showed similar proportions of dead pollen (Figure 2).

**Table 2:** Segregation analyses at L3 of selfed progenies in two different nuclear backgrounds

| Genotype | Number of plants <sup>(1)</sup> | | | | $p \chi^2 (1:2:1)$ | $f \text{Sha}^{(2)}$ |
| --- | --- | --- | --- | --- | --- | --- |
|  | Sha | Hz | Mr-0 | Total |  |  |
| MrL3<sup>H</sup> | 14 | 83 | 73 | 170 | $1.2 \cdot 10^{-9} ***$ | 0.08 |
| ShaL3<sup>H</sup> <sup>(3)</sup> | 18 | 87 | 73 | 178 | $4.0 \cdot 10^{-8} ***$ | 0.10 |

<sup>(1)</sup> genotyped at marker M1 (described in Figure 5 Source Data 3). <sup>(2)</sup> Frequency of Sha homozygotes (expected frequency 0.25). <sup>(3)</sup> from Simon *et al.* (2016). \*\*\* $p < 0.001$ .

**Figure 2:**
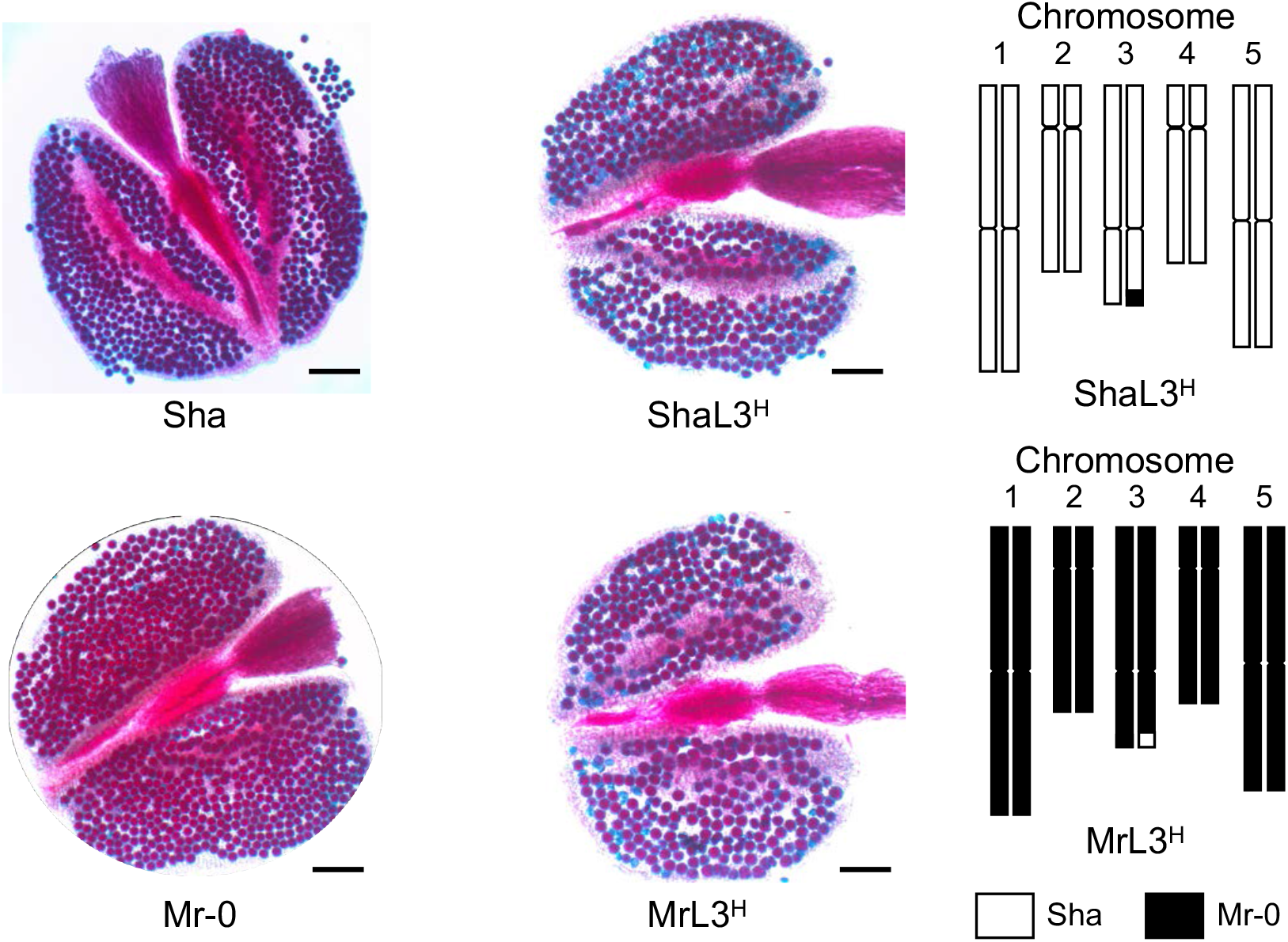
Plants heterozygous at L3 in either Sha or Mr-0 nuclear backgrounds show dead pollen. Typical pollen phenotypes (Alexander staining of anthers) of ShaL3^H^ and MrL3^H^ plants compared to the parental accessions Sha and Mr-0. Viable pollen grains are stained in red, aborted pollen grains appear in blue. Scale bars: 100µm. Graphical representations of ShaL3^H^ and MrL3^H^ genotypes are displayed on the right.

We further studied the ShaL3^H^ genotype, which flowers earlier than MrL3^H^. We observed a strong distortion when ShaL3^H^ was used as male in a cross with Sha, but no distortion when it served as female parent (Table 3). These results confirmed that the segregation distortion was only due to a PK.

**Table 3:**
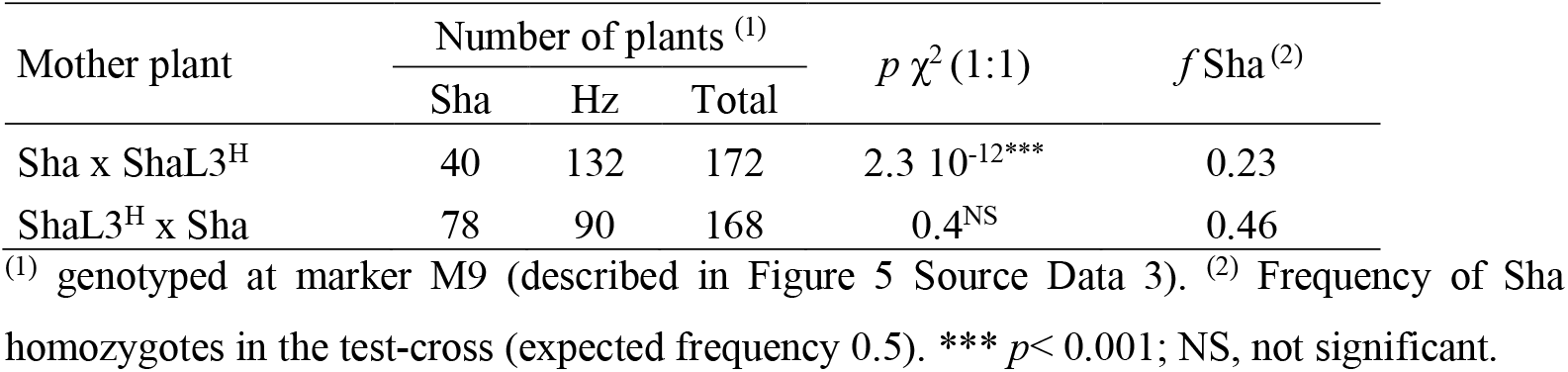
Transmission of the Sha allele in a heterozygous context from the male and female sides

We used cytological approaches to determine whether the male dysfunction occurred during meiosis or during pollen development and to specify its timing. The male meiosis of ShaL3^H^ plants was identical to that of fixed siblings at all stages (Figure 3), excluding a meiotic defect as the source of the segregation bias in ShaL3^H^ plants. During male gametogenesis, abnormal pollen grains were observed in anthers of ShaL3^H^ plants from the bicellular stage, after the first pollen division (Figure 4). The proportion of abnormal pollen increased in the tricellular stage, and about 35% of mature pollen grains were dead in ShaL3^H^ plants (Figure 4), which reflects an incomplete penetrance since 50% dead pollen would be expected if the PK effect was total. We concluded that, in these plants, the Sha allele at L3 is poorly transmitted because most of the Sha pollen grains fail to develop properly from the binucleate stage and eventually die.

**Figure 3:**
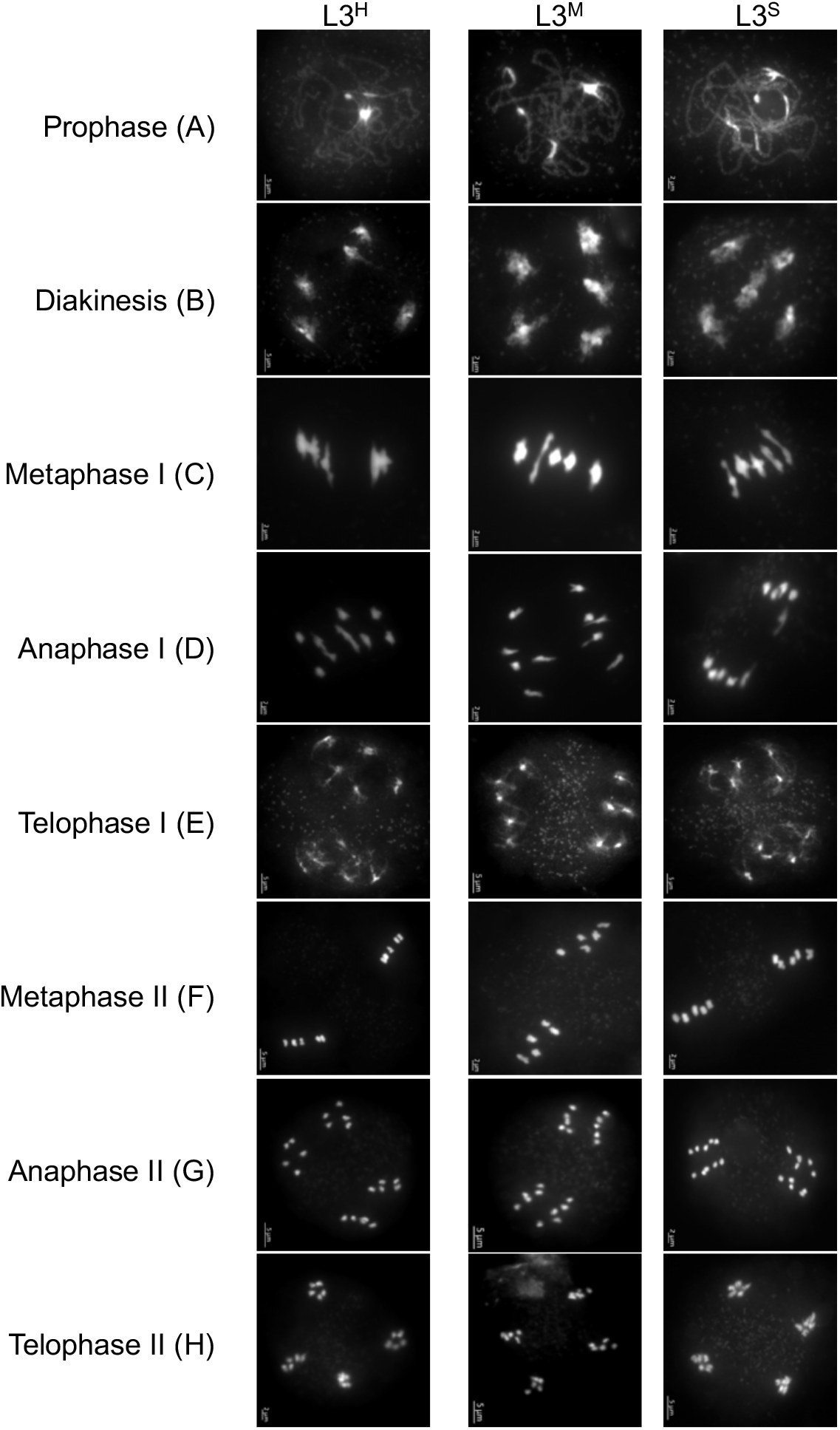
Plants heterozygous at L3 in Sha nuclear background have a normal male meiosis. The meiotic progression was analyzed by DAPI staining of meiotic chromosome spreads on anthers from siblings from a ShaL3^H^ plant. During prophase I (A), meiotic chromosomes condense, recombine and undergo synapsis, resulting in the formation of five bivalents which become visible at diakinesis (B). The bivalents align at metaphase I (C), and chromosomes separate from their homologues at anaphase I (D), leading to the formation of two pools of five chromosomes and two nuclei (E). At the second meiotic division, the pairs of sister chromatids align on the two metaphase plates (F), and separate at anaphase II (G) to generate four pools of five chromosomes, which give rise to tetrads of four microspores (H). The meiotic progression in plants heterozygous at L3 (L3^H^, N=322) is identical to those homozygous Sha (L3^S^, N=166). and Mr-0 (L3^M^, N=77).

**Figure 4:**
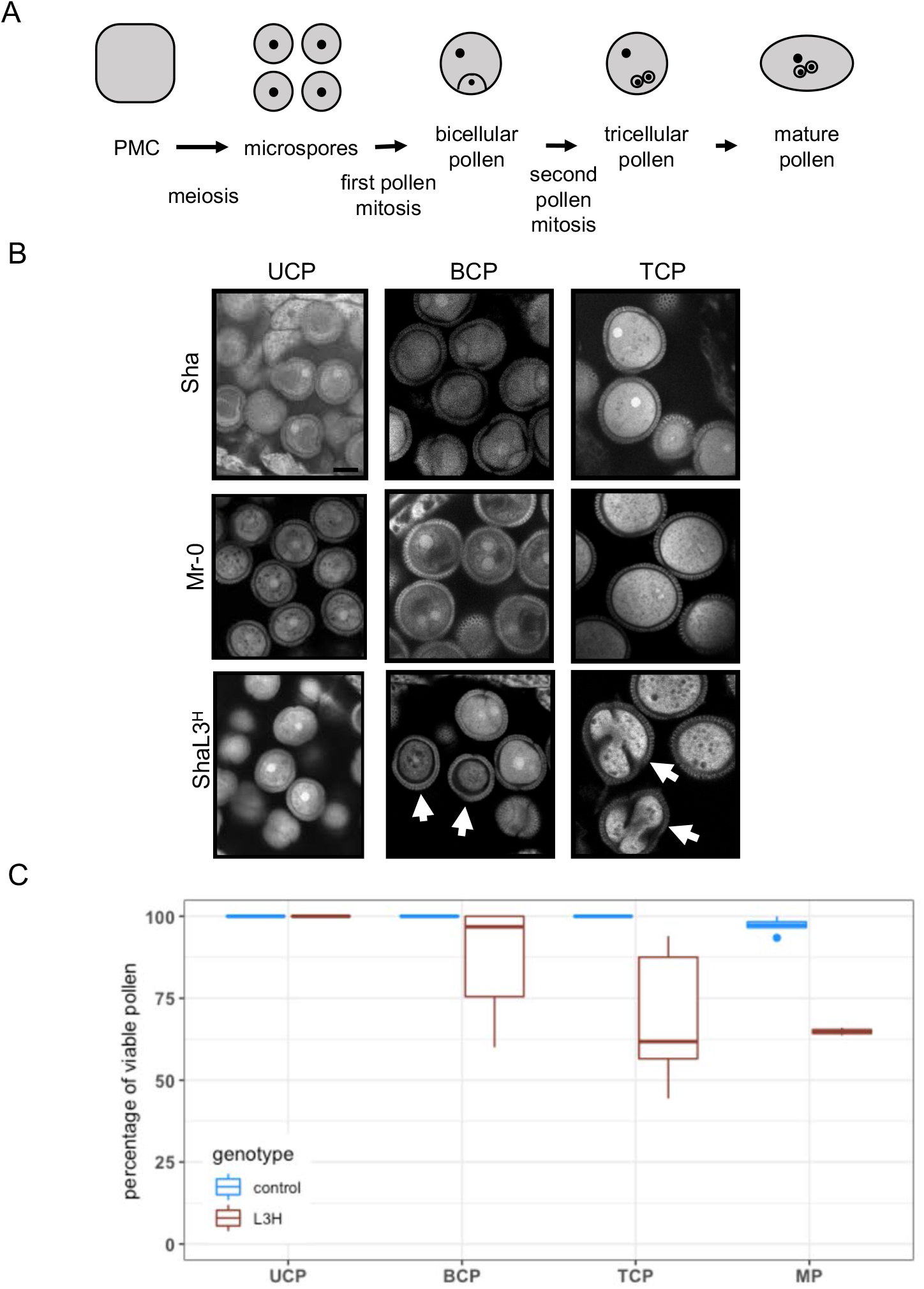
Pollen from plants heterozygous at L3 abort progressively from bicellular stage. A: The scheme outlines the main steps of pollen development in *A. thaliana*: the pollen mother cell (PMC) produces haploid microspores (or unicellular pollen, UCP) through meiosis; the first pollen mitosis produces bicellular pollen (BCP); the reproductive cell divides in the second pollen mitosis to give tricellular pollen (TCP), which evolves into mature pollen (MP). B: Typical confocal images of male gametogenesis in ShaL3^H^ plants (bottom row) and in Sha and Mr-0 parental controls after propidium iodide (PI) staining. White arrows indicate abortive pollen. Scale bar: 10µm. C: Percentage of normal pollen (Figure 4-Source Data 1) in anthers of pooled parental (blue) and ShaL3^H^ (brown) plants at different stages of development (countings from confocal acquisitions of PI stainings), and in mature pollen (countings from Alexander staining).

### The PK at L3 contains three genetic elements

L3 was previously mapped in a 280 Kb interval at the bottom of chromosome 3 (Simon et al. 2016). We fine-mapped the PK in the genotype ShaL3^H^ using the presence of a segregation bias in the self-descent of recombinants as a robust phenotypic trait to narrow down the L3 interval: genetic markers fixed for Sha or Mr-0 alleles in recombinants with a significant bias in their progeny were excluded from the candidate interval. This strategy allowed us to map all the genetic elements necessary for the PK activity in an interval hereafter called PK3, corresponding to the region flanked by markers M5 and M13 (68 Kb in Col-0) (Figure 5A). Out of a total of 4,717 plants genotyped, we found six recombinants between M5 and M13. Recombination points of these recombinants, 27D6, 25A7, 52D12, 52D7, 8F10BH2 and 23G9 were finely localized. Only three of these plants, 52D7, 8F10BH2 and 23G9, were recombined between M6 and M12, which are very close to M5 and M13, respectively (Figure 5 Source Data 1). None of them presented a bias in its progeny whereas 27D6, 25A7 and 52D12, which are heterozygous between M6 and M12, did. Pollen viability of the recombinants, assessed by Alexander stainings, were consistent with the presence or absence of bias in their progenies (Figure 5B). Further information on the genetic structure of the PK3 was obtained from the three plants recombined between M6 and M12. First, by crossing 52D7 and 8F10BH2 with Sha, we converted their fixed portion of the interval into a heterozygous region (Figure 5A). The offsprings of these new genotypes (named i-52D7 and i-8F10BH2) did not show any segregation bias, whereas their siblings heterozygous along the whole interval did (Figure 5 Source Data 2). This indicated that the parts of the PK3 interval that were heterozygous in 52D7 and in 8F10BH2 both contained elements necessary for the PK activity. We thus segmented the PK3 locus into three genetic intervals, named PK3A, PK3B and PK3C (Figure 5A), PK3A and PK3C both carrying elements necessary for a functional PK. Then, in order to evaluate the PK3B interval, we crossed fixed progenies of 23G9 and 52D7 that inherited the recombination events from their parents, and obtain a plant (23G9#15 × 52D7#7) heterozygous at both PK3A and PK3C and fixed Mr-0 at PK3B (Figure 5A). The absence of segregation bias in the selfed progeny of this plant showed that PK3B also carried an element necessary for the PK activity. Therefore, each of the three parts of the PK3 interval contains at least one element required for the PK activity. On one hand, Mr-0 alleles were required at PK3A and at PK3C: these two intervals thus contain killer elements. On the other hand, when the Sha allele was absent at PK3B while PK3A and PK3C were heterozygous, the PK was no longer active, indicating that either a target element from Sha was missing, or an antidote from Mr-0 was present in all the pollen grains produced.

**Figure 5:**
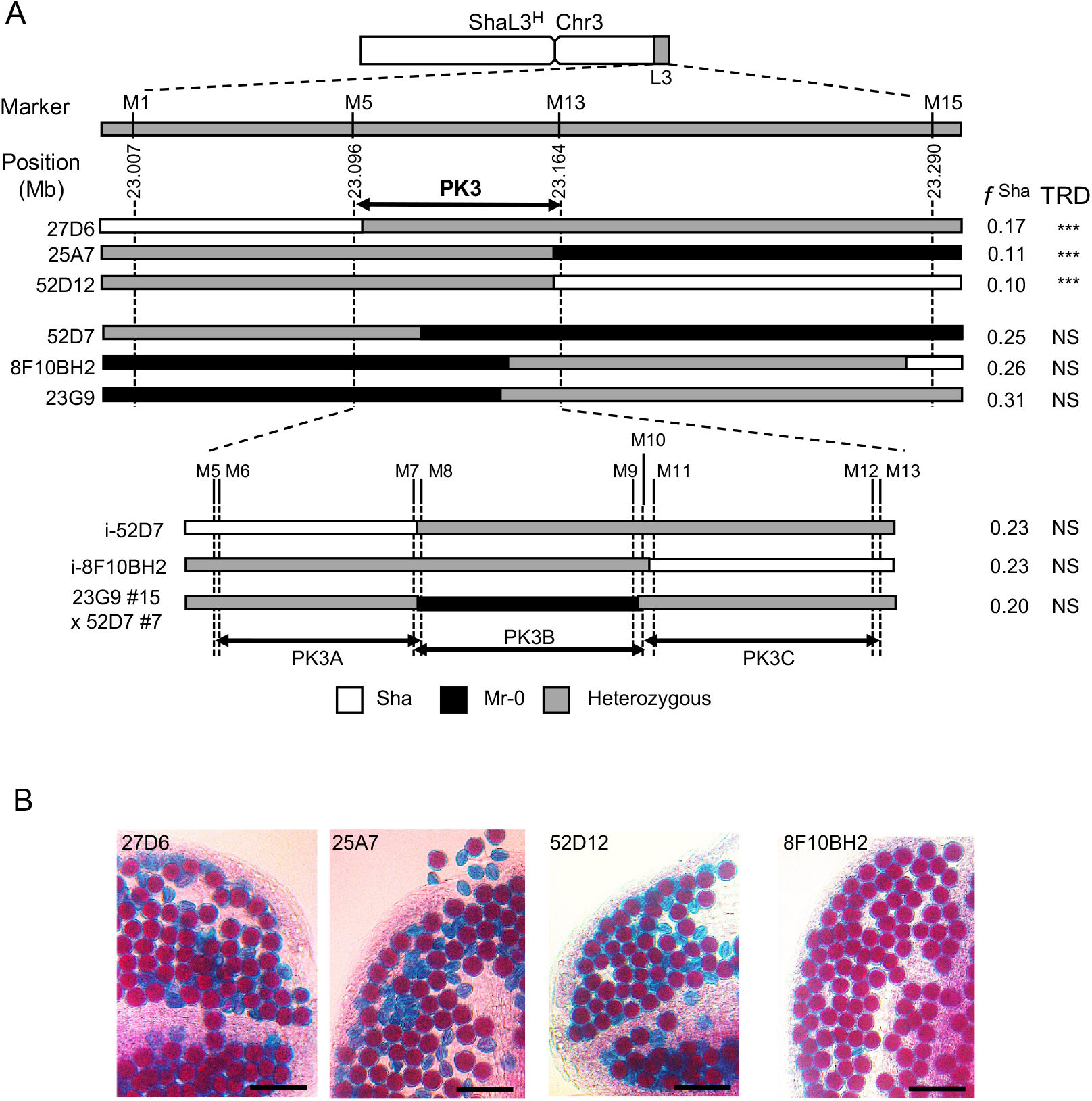
Fine mapping of the PK3 locus. A: The upper panel shows the mapping of PK3 in the L3 locus delimited by the genetic markers M1 and M15 (Figure 5-Source Data 1). Markers M5 and M13 delimit the PK3 interval, where all the PK elements required to cause the segregation bias are present. Positions of markers on the TAIR10 Col-0 genomic sequence are given. The most relevant recombinants are shown, with the frequencies of Sha homozygotes (*f* ^Sha^) observed in their selfed progenies indicated on the right. TRD, transmission ratio distortion (chi square test for 1:2:1 segregation) *** *p* < 0.001, NS: not significant. The lower panel shows the further dissection of the PK3 locus into PK3A, PK3B and PK3C (Figure 5-Source Data 2). Because some markers are very close to each other (Figure 5-Source Data 1), this diagram is not to scale. Genotypes used to dissect the PK3 are presented: i-52D7 and i-8F10BH2 result from crosses of 52D7 and 8F10BH2, respectively, with Sha; the 23G9#15 × 52D7#7 genotype, with two heterozygous regions flanking a fixed central region of the PK3, was produced by crossing appropriate selfed progenies of 23G9 and 52D7 recombinants. All markers are described in Figure 5-Source Data 3. B: Pollen viability in recombinants used to delineate the PK3 interval. Representative images of Alexander staining of anthers from recombinants presented in A. Aborted pollen grains are blue whereas living pollen grains are red. Scale bar: 50µm.

### The PK3 locus is highly variable

To highlight differences between Sha and Mr-0, we sequenced the entire locus in both accessions. The overall structure of the PK3 locus in Sha was very similar to that of Col-0 (Figure 6A), the main differences being the deletion of the transposable element (TE) *AT3G62455*, a 1308 bp insertion in the intron of *AT3G62460*, and an insertion of approximately 1 Kb in the intergenic region between *AT3G62540* and *AT3G62550*. In contrast, the PK3 locus in Mr-0 locus was particularly complex as compared to Col-0 and Sha, showing many structural variations such as large deletions, insertions, duplications and inversions (Figure 6A). Two TEs present in Col-0, *AT3G62455* and *AT3G62520*, were missing in Mr-0. The TEs *AT3G62475, AT3G62480* and *AT3G62490* that are located in the PK3A region of Sha, were absent from this region in Mr-0, but its PK3B region presented a large insertion of over 20 Kb that contained several TEs including *AT3G62475, AT3G62480* and *AT3G62490* homologues. Nonetheless, the same protein coding genes are present in the three genotypes, even though Mr-0 has two copies of the gene *AT3G62510* and three copies of the genes *AT3G62528, AT3G62530* and *AT3G62540*, with one copy of *AT3G62530* and *AT3G62540* being inserted into the second intron of *AT3G62610* (Figure 6A).

**Figure 6:**
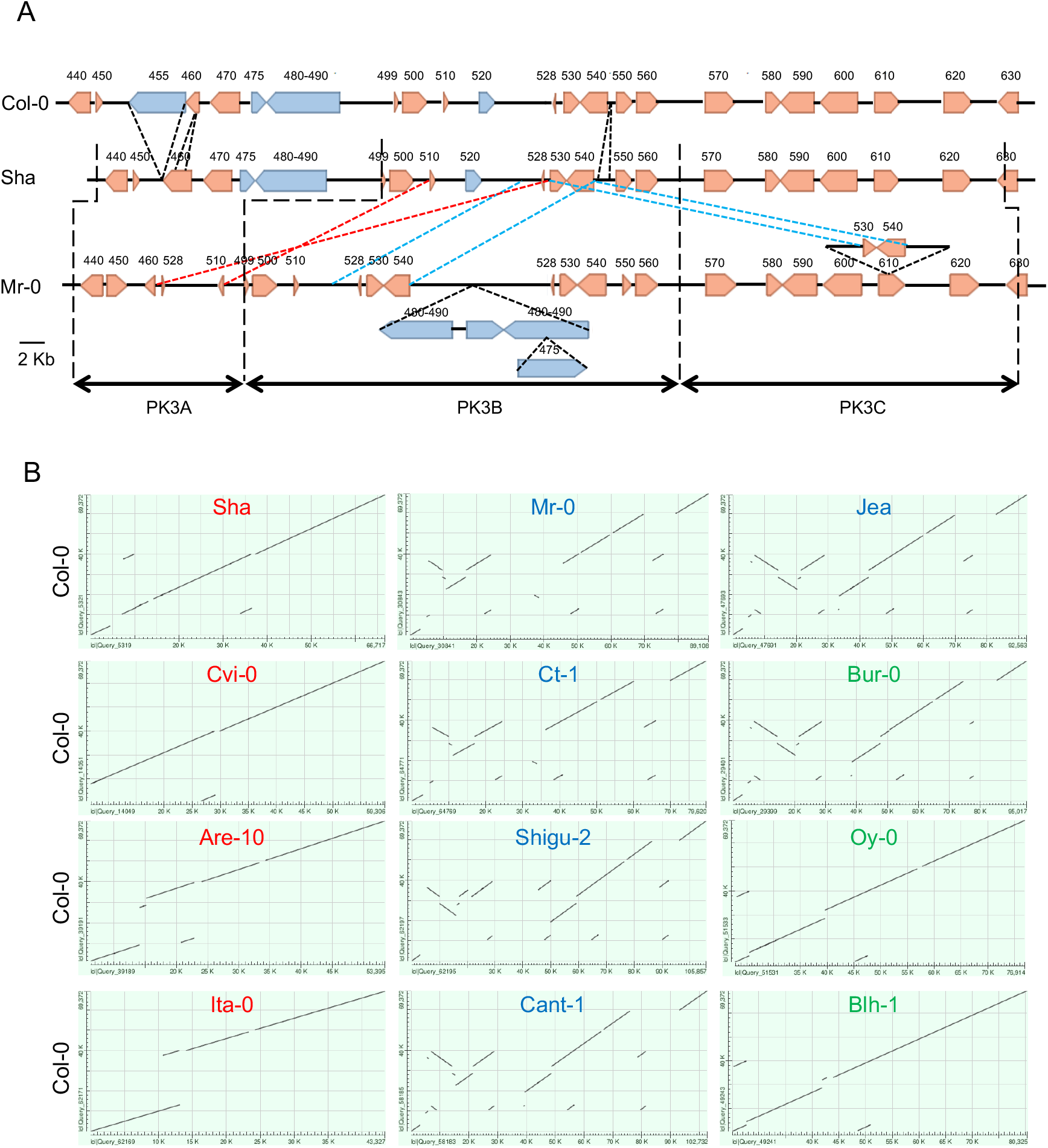
Structural variation at the PK3 locus. A: PK3 locus in Col-0, Sha and Mr-0. Beige and blue plain arrows represent protein coding genes and TEs, respectively, with their orientations. Gene labels on the Col-0 locus correspond to the last three digits of AT3G62xxx gene identifiers in the reference sequence (TAIR10), the same numbers on Sha and Mr-0 loci indicate genes homologous to the Col-0 genes. Black, red and blue dotted lines respectively represent insertions/deletions, inversions and duplications. The limits of the PK3A, PK3B and PK3C intervals in Sha and Mr-0 are indicated (dashed broken lines). B: Dot plots of PK3 sequences of 12 accessions (x axes) against the reference Col-0 (y axes), generated at NCBI using the nucleotide Blast tool with the option ‘align two or more sequences’ and the following settings: max target = 10, Expect threshold = 0.001, word size = 256 (Altschul et al. 1990). The names of killer, neutral and killed accessions are in blue, green and red, respectively. The sequences can be accessed by DOI numbers indicated in Figure 6 Source Data 1, with positions of the PK3 sequences.

Because the PK3 locus is highly rearranged between these three accessions, we looked at other variants of known status for the PK phenotype. The entire genomes of 10 such variants, four killers, three killed and three neutral, were *de novo* sequenced. PK3 sequence alignments revealed structural variations relative to Col-0 in all the killers (Figure 6B). On the contrary, PK3 loci of killed and neutral natural variants, excepted Bur-0, are mostly colinear with Col-0 (Figure 6B). When comparing the synteny of protein coding genes at the locus between *A. thaliana* accessions and *A. lyrata* (Figure 7), we observed that the locus structure in Ita-0 is similar to that of *A. lyrata*. This locus structure is also found in other Brassicaceae, such *Boechera stricta* (Figure 7). In contrast, Col-0, the two other neutral accessions Blh-1 and Oy-0, and Sha have a duplication of the *A. lyrata* gene *AL5G45290*, that encodes a pentatricopeptide repeat protein (PPR), resulting in two nearly identical genes (*AT3G62470* and *AT3G62540*). Five additional protein coding genes (*AT3G62499* to *AT3G62530*) compared to the *A. lyrata* and Ita-0 sequences were found between the two PPR paralogues in these accessions. We also observed a great variability in the number of copies of all these genes according to the accessions. Compared to Col-0, Blh-1, Oy-0 and Sha, where they were present once, some of them were absent in the other killed accessions Cvi-0 and Are-10. On the contrary, in the killers, they have undergone a variable number of duplications, with one to four copies depending on both the genes and the accessions (Figure 7).

**Figure 7:**
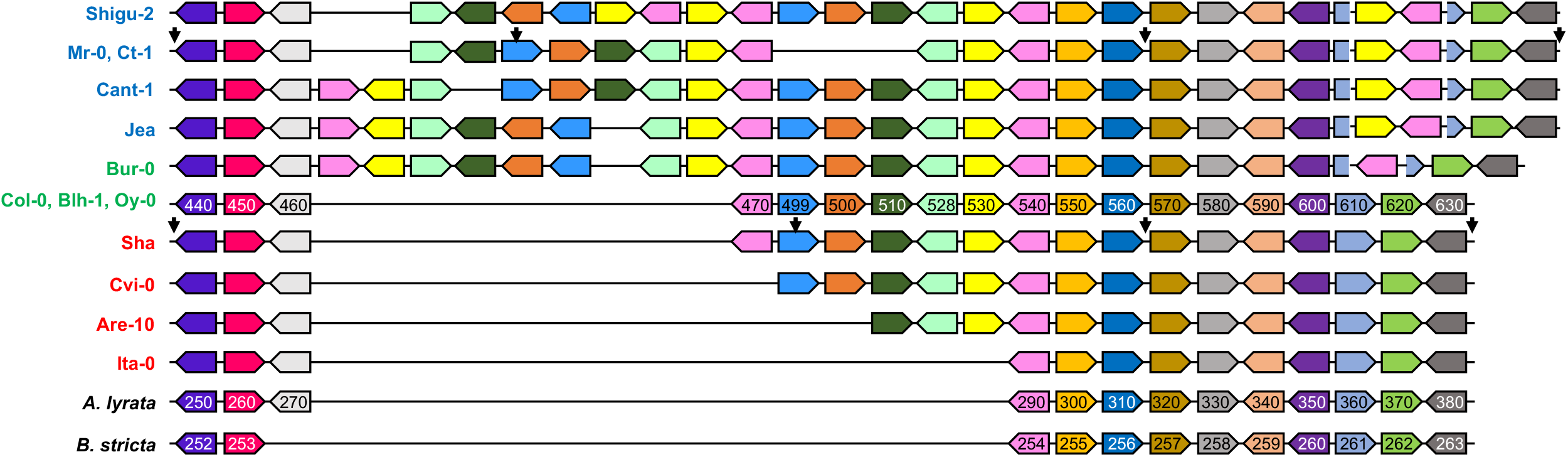
Intra and Inter-specific synteny of protein coding genes at the PK3 locus. Alignment the PK3 loci from thirteen *A. thaliana* accessions and two related species, *Arabidopsis lyrata* and *Boechera stricta*, drawn to highlight synteny between protein coding genes. The scheme is not to scale and TEs are not represented. For *A. lyrata* and *B. stricta* the structural annotation was obtained from Phytozome (https://phytozome.jgi.doe.gov/). The names of killer, neutral and killed *A. thaliana* accessions are written in blue, green and red, respectively. Plain coloured arrows represent coding genes with their orientations, each colour representing orthologues and paralogues of a same gene. Gene labels correspond to the last three digits of AT3G62xxx, AL5G45xxx and Bostr.13158s0xxx gene identifiers in *A. thaliana* (Col-0 reference sequence TAIR10), *A. lyrata* (V2.1) and *B. stricta* (V1.2), respectively. Black arrows above the Mr-0 and Sha loci delimit the PK3A, PK3B and PK3C intervals from left to right.

Given the complexity and diversity of the locus, no obvious candidate gene appeared for killer or target/antidote elements. For the killer elements, an additional difficulty results from the fact that two elements are necessary for the killing activity, one located in the PK3A interval and the other in the PK3C interval, thereby preventing to draw conclusions from the comparison of killer and non-killer alleles in a single interval. Moreover, the intervals PK3A, PK3B and PK3C were delimited by the mapping recombinants between Sha and Mr-0, and, due to the structural differences between the accessions, it is not possible to infer their frontiers in the other genotypes, at least for the limit between PK3A and PK3B. However, it can be noted that the locus of the neutral accession Bur-0 is nearly identical to that of the killer Jea: only the copy of *AT3G62530* inserted into *AT3G62610* in the PK3C interval in all the killers is missing in Bur-0 (Figure 7). This makes this copy, which is absent in all the non-killer alleles, a gene to be tested for killer activity, although it cannot be excluded that Bur-0 lost its killer activity due to another polymorphism in PK3C or in its unknown PK3A killer element.

We thus focused on the PK3B interval, which contains either a target element in killed alleles or an antidote in killer and neutral alleles. However, comparison of the PK3B sequences did not reveal a gene specific for the killed alleles that could encode a target, nor a gene specific for the killer and neutral alleles that could encode an antidote. We therefore exploited mutants in each of the genes present in this region.

### *AT3G62530*, expressed in young developing pollen, encodes the antidote

Col-0 had a neutral behaviour regarding the PK (Table 1). That means that, in a poison-antidote model, Col-0 would carry an antidote element whose inactivation should let Col-0 pollen unprotected against Mr-0 killer. A hybrid between Col-0 with an inactivated version of the antidote and Mr-0 would then present dead pollen, and produce an F2 with a segregation bias against the Col-0 allele at L3. In order to test the poison-antidote model, we thus used T-DNA insertion mutants available in this genetic background for the eight PK3B genes. None of the homozygous mutants (Col^mut^) was distinguishable at the phenotypic level from Col-0 in our greenhouse conditions. We crossed each Col^mut^ with an early-flowering Mr-0 genotype carrying a KO mutation in the *FRIGIDA* gene, hereafter named Mr*fri* (see Materials and Methods). All the Mr*fri* x Col^mut^ F1 plants exhibited only viable pollen, except the hybrid with a T-DNA insertion in *AT3G62530* (Col^mut530^) that presented aborted pollen (Figure 8). Then, we genotyped the F2 families at L3, which all followed the expected Mendelian proportions, with the exception of the Mr*fri* x Col^mut530^ and Mr*fri* x Col^mut499^ F2s. The latter presented a slightly biased segregation against the Col^mut499^ allele, but when the Mrfri x Col^mut499^ F1 was crossed as female or male parent using Mr*fri* as a tester, no mutant allele transmission bias was detected (Figure 8). In contrast, the Mr*fri* x Col^mut530^ F2 population showed a strong bias against Col^mut530^ homozygous genotypes, as those observed in hybrids with an active PK. We analysed the mutant allele transmission in the presence of a killer allele by crossing the Mr*fri* x Col^mut530^ F1 as female or male with Mr*fri* as a tester, and we observed a significant bias against the Col^mut530^ allele only when it came from pollen (Figure 8). At this step, we characterized the T-DNA insertion in *AT3G62530*, we checked that it really inactivated the gene and did not induce pollen abortion nor a bias in the transmission of the mutation to the progeny of the heterozygous Col^mut530^ mutant (Table 4). These results strongly suggested that *AT3G62530* encoded an antidote. If this is the case, the mutation in *AT3G62530* should have no effect on pollen viability nor on allele segregation in the absence of a killer allele. Indeed, no dead pollen was observed in the anthers of homozygous nor heterozygous Col^mut530^ (Figure 8 Supplement 2), and no segregation bias against the Col^mut530^ allele was detected in the self-progeny of the heterozygous Col^mut530^ nor in the progeny of the Sha x Col^mut530^ hybrid (Table 4). We concluded that the presence of a killer allele was necessary to trigger the death of Col^mut530^ pollen. This was confirmed by crossing Col^mut530^ with another killer accession, Ct-1: while no bias was found in the progeny of the Ct-1 x Col-0 hybrid, a bias against the Col^mut530^ allele was observed in the progeny of Ct-1 x Col^mut530^ (Table 4), which was very similar to the bias against the Sha allele observed in the progeny of the Ct-1 x Sha cross.

**Figure 8:**
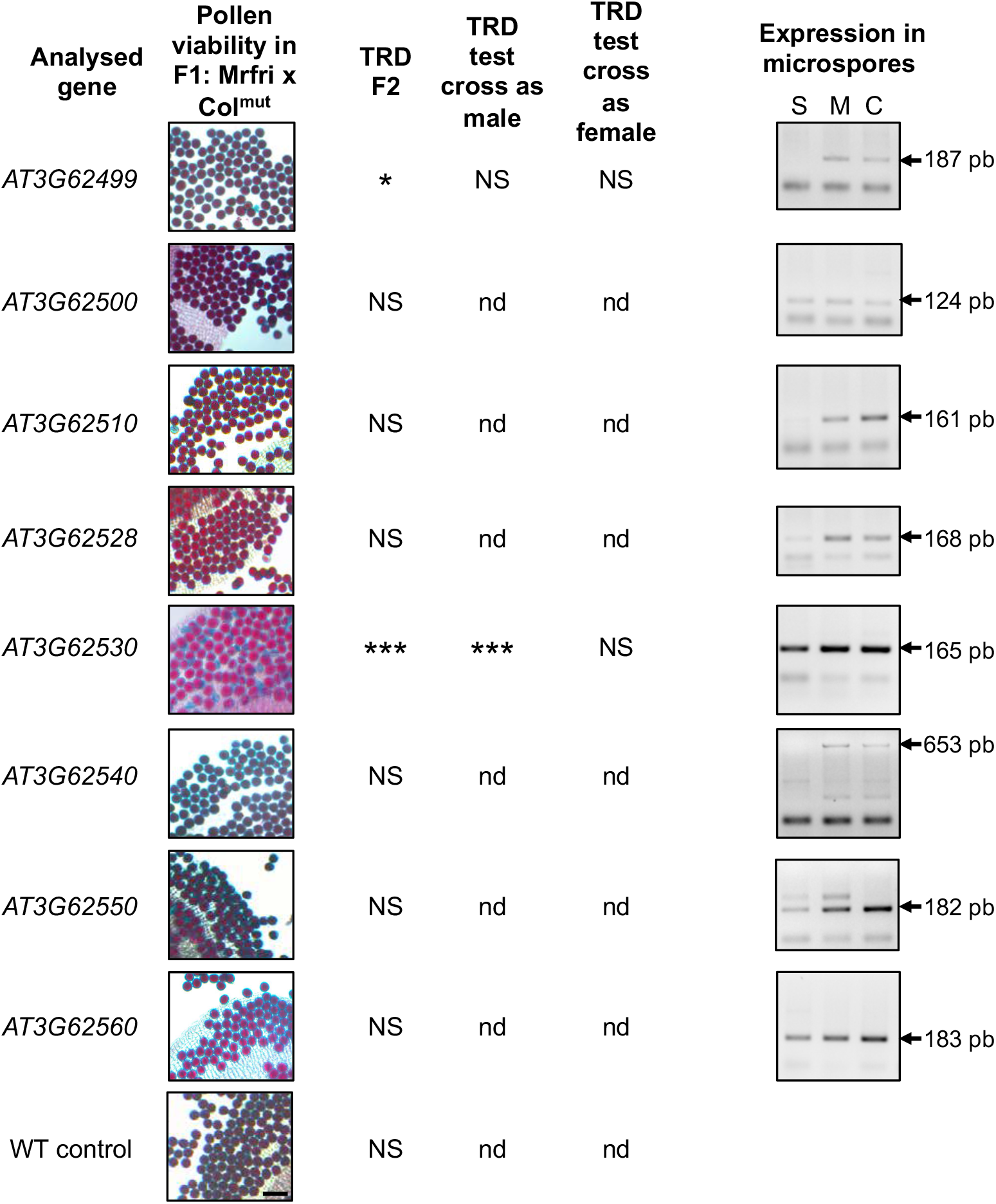
Analysis of mutants and microspore expression for PK3B coding genes. We tested each coding gene at PK3B for an putative antidote behavior. Each homozygous Col-0 mutant (Figure 8-Source Data 1) was crossed with Mr*fri* (Figure 8 Supplement 1) and we observed pollen viability (Alexander staining of anthers) in each Mr*fri* x Col^mut^ F1. Viable pollen grains are stained in red, aborted pollen grains appear in blue. Scale bar: 50µm. F2 progenies of each F1 were tested for a transmission ratio distortion (TRD) at PK3 (Figure 8 – Source Data 2). The transmission of Col^mut499^ and Col^mut530^ alleles through male and female gametes were tested in test crosses (Figure 8-Source Data 3). NS, not significant; * pvalue <0.05; ***pvalue < 0.001; nd, not determined. The T-DNA insertion in the Colmut^530^ allele was characterized (Figure 8 Supplement 2). On the right, RT-PCR results for expression of PK3B genes from Sha (S), Mr-0 (M) and Col-0 (C) alleles in purified microspores are given (Figure 8 – Source Data 4). The Mr-0 allele was analysed in a progeny of Sha L3^H^ homozygous Mr-0 at L3. PCR primers used (Supplemental File 1) do not allow to distinguish the two or three copies of *AT3G62510, AT3G62528, AT3G62530* and *AT3G62540* that exist in the Mr-0 allele. The expected sizes of the amplification products are given on the right.

**Table 4:** Segregation at L3 in F2s where the Col^mut530^ allele is confronted to a killed (Sha), neutral (Col-0) or killer (Ct-1) allele

| Genotype | Number of plants | | | | $p \chi^2 (1:2:1)$ | $f \text{Col}^{\text{mut530}} (4)$ |
| --- | --- | --- | --- | --- | --- | --- |
|  | not Col<sup>mut530</sup> | Hz | Col<sup>mut530</sup> | Total |  |  |
| F2 (Sha x Col<sup>mut530</sup>)<sup>(1)</sup> | 46 | 101 | 37 | 184 | 0.3<sup>NS</sup> | 0.20 |
| heterozygote Col<sup>mut530</sup>/Col-0<sup>(2)</sup> | 49 | 85 | 46 | 180 | 0.5<sup>NS</sup> | 0.26 |
| F2 (Ct-1 x Col<sup>mut530</sup>)<sup>(3)</sup> | 74 | 89 | 21 | 184 | 2 10<sup>-7</sup> *** | 0.11 |

<sup>(1)</sup> genotyped with marker M1. <sup>(2)</sup> genotyped with PCR primers used to characterise the mutant (Figure 8 Supplement 2). <sup>(3)</sup> The progeny of the F1 (Ct-1 x Col-0) presents no TRD (Table 4 Supplement 1). <sup>(4)</sup> frequency of Col<sup>mut530</sup> homozygotes (expected frequency 0.25). \*\*\* p < 0.001; NS, not significant.

In the frame of the poison-antidote model, the antidote must be expressed in cells that need to be protected from the poison elements, in particular in developing pollen. Indeed, RT-PCR assays showed that *AT3G62530*, in addition to being expressed in leaves, is expressed in microspores (young developing pollen, before the first pollen mitosis, Figure 4) from plants either Col-0, Sha or Mr-0 at the locus (Figure 8). In addition, *AT3G62530* seems one of the most expressed gene in microspores amongst those of the PK3B interval, in the three genotypes. All together, the above results fit perfectly with a poison-antidote system where *AT3G62530* codes the antidote, Col-0 and Mr-0 carrying functional forms of the antidote while Sha has a non-functional antidote allele. We named the gene *APOK3*, for ***A****NTIDOTE OF* ***PO****LLEN* ***K****ILLER ON CHROMOSOME* ***3***.

### *APOK3* encodes a chimeric protein addressed to mitochondria

Because APOK3 is expressed at similar levels in microspores carrying the Sha allele, not protected by an antidote, and microspores carrying Col-0 or Mr-0 alleles, which must harbor an active antidote (Figure 8), we hypothesized that the allelic differences for antidote activity are due to differences at the protein level.

In Col-0, *APOK3* encodes a protein of 221 amino acids, belonging to the ARM-repeat superfamily (https://www.arabidopsis.org/), defined by the presence of tandem repeats generally forming alpha-helices (https://supfam.org/). Further analysis of structural domains identified three HEAT repeat domains (Figure 9A). Different parts of APOK3 are very similar to parts of proteins encoded by the genes *AT3G62460* (98% identity on residues 1 to 44), *AT3G43260* (66% identity and 73% similarity on residues 43 to 142) and *AT3G58180* (57% identity and 78% similarity on residues 81 to 221) (Figure 9B). These three genes are located on the chromosome 3, and it is interesting to note that *AT3G62460* is in the PK3A interval. AT3G43260 and AT3G58180 are both annotated as related to deoxyhypusine hydroxylases of other organisms, but a close examination showed that the genuine Col-0 deoxyhypusine hydroxylase is encoded by *AT3G58180* (Figure 9 Supplement 1). The part of the AT3G62460 protein shared by APOK3 includes a mitochondria-targeting peptide (Figure 9A), suggesting that APOK3 is addressed to mitochondria. Indeed, APOK3 has been repeatedly found in *A. thaliana* mitochondrial proteomes (Heazlewood et al. 2004; Klodmann et al. 2011; Taylor et al. 2011; Konig et al. 2014; Senkler et al. 2017). We therefore hypothesized that, if the antidote acts in the mitochondria, the strength of the bias due to the PK could be influenced by the mother plant cytoplasmic background. As a first insight in this direction, we compared the segregation distortion in the progenies of plants heterozygous Sha/Mr-0 at PK3 differing only by their cytoplasmic backgrounds, and we observed that the bias was stronger in the Sha than in the Mr-0 cytoplasmic background (Figure 9C). We concluded that the antidote function of APOK3 is likely to be sensitive to variation in the mitochondrial genome.

**Figure 9:**
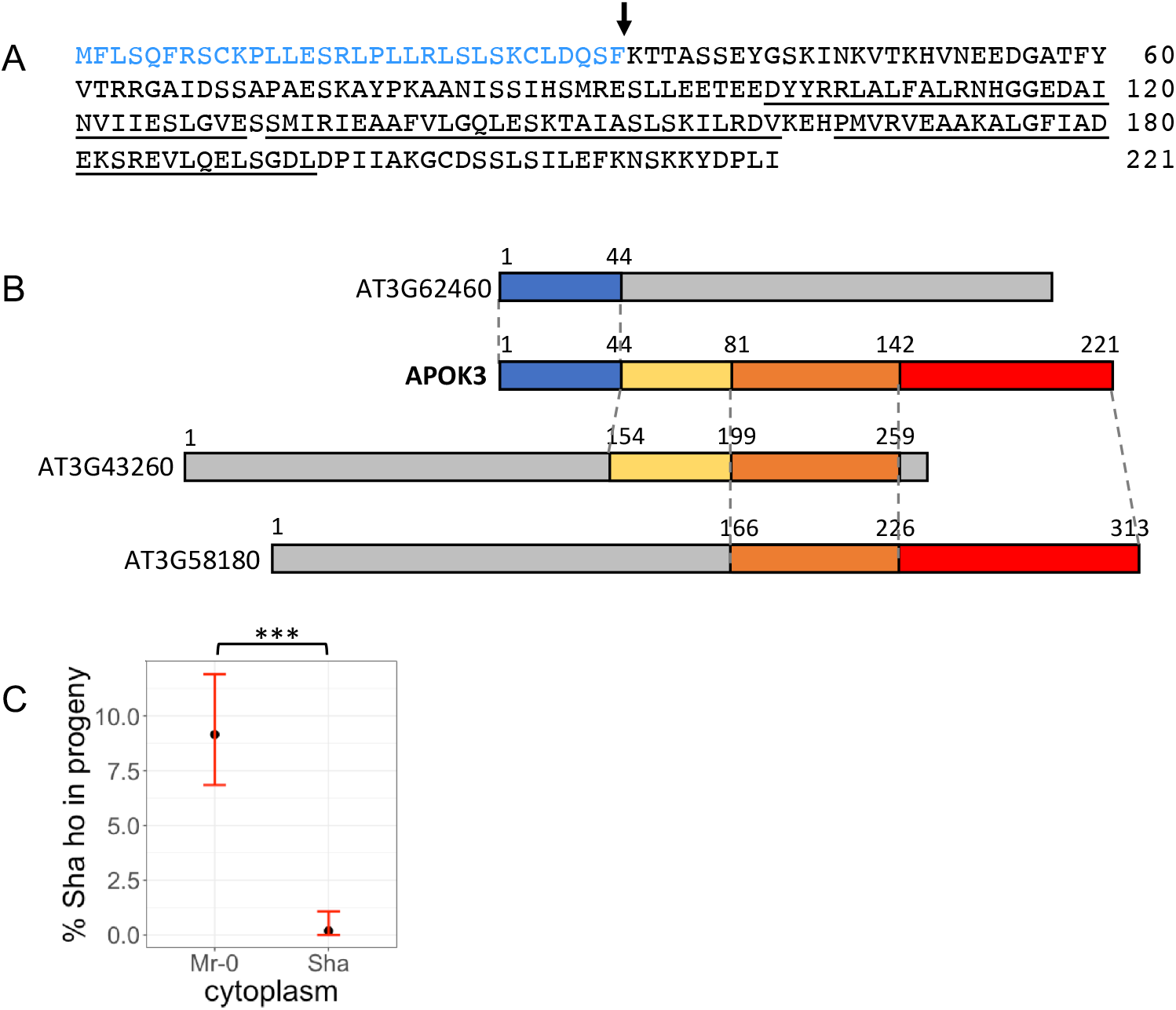
Structure of APOK3. A: Protein sequence of APOK3 in Col-0. The N-terminal mitochondria-targeting peptide (in blue) with a potential cleavage site between amino acids 32 and 33 (arrow) was predicted by TargetP2.0 (likelihood = 0.9; http://www.cbs.dtu.dk/services/TargetP/) (Almagro Armenteros et al. 2019). The three HEAT-repeat domains (underlined) were identified with SMART (Letunic et al. 2021). B: Representation of the composite APOK3 structure. Colours indicate the homologous regions between APOK3 and AT3G62460 (blue), AT3G43260 (yellow) and AT3G58180 (*A. thaliana* deoxyhypusine hydroxylase, Figure 9-Supplement 1) (red). The orange box corresponds to a region homologous between APOK3 and both AT3G43260 and AT3G58180. Grey boxes indicate regions of the proteins not aligning with APOK3. Genes encoding proteins similar to APOK3 in the Col-0 genome were retrieved through BlastP search (2.9.0+ default settings, Altschul et al, 1997) against Araport11 protein sequences on TAIR. C: Effect of cytoplasmic background on the strength of the PK3 induced TRD. Percentages of Sha homozygotes at PK3 were measured in three independent progeny pairs of reciprocal F1s that display the Sha/Mr-0 PK3 phenotype, in either Mr-0 or Sha cytoplasmic background. They were obtained by crossing both way with Sha three mapping recombinants and selecting F1s that were heterozygous on the whole PK3 interval (Figure 9 Source data 1). Dots indicate the average of Sha homozygote percentage for each cytoplasmic background. Red vertical bars indicate 0.95 confidence intervals. *** Fisher test *p* < 0.001.

### *APOK3* has undergone several duplication events within killer PK3 loci

Mr-0 has two strictly identical copies of *APOK3* in the PK3B interval, which have the same structure as Col-0 and Sha genes. Mr-0 also has a third copy inserted with other sequences in the intron of *AT3G62610* in the PK3C interval (Figure 10A), but the N-terminal part of this copy differs from that of the other two and is not homologous to *AT3G62460*, which suggests that this copy is functionally different from the others; it was thus named *APOK3-like*.

**Figure 10:**
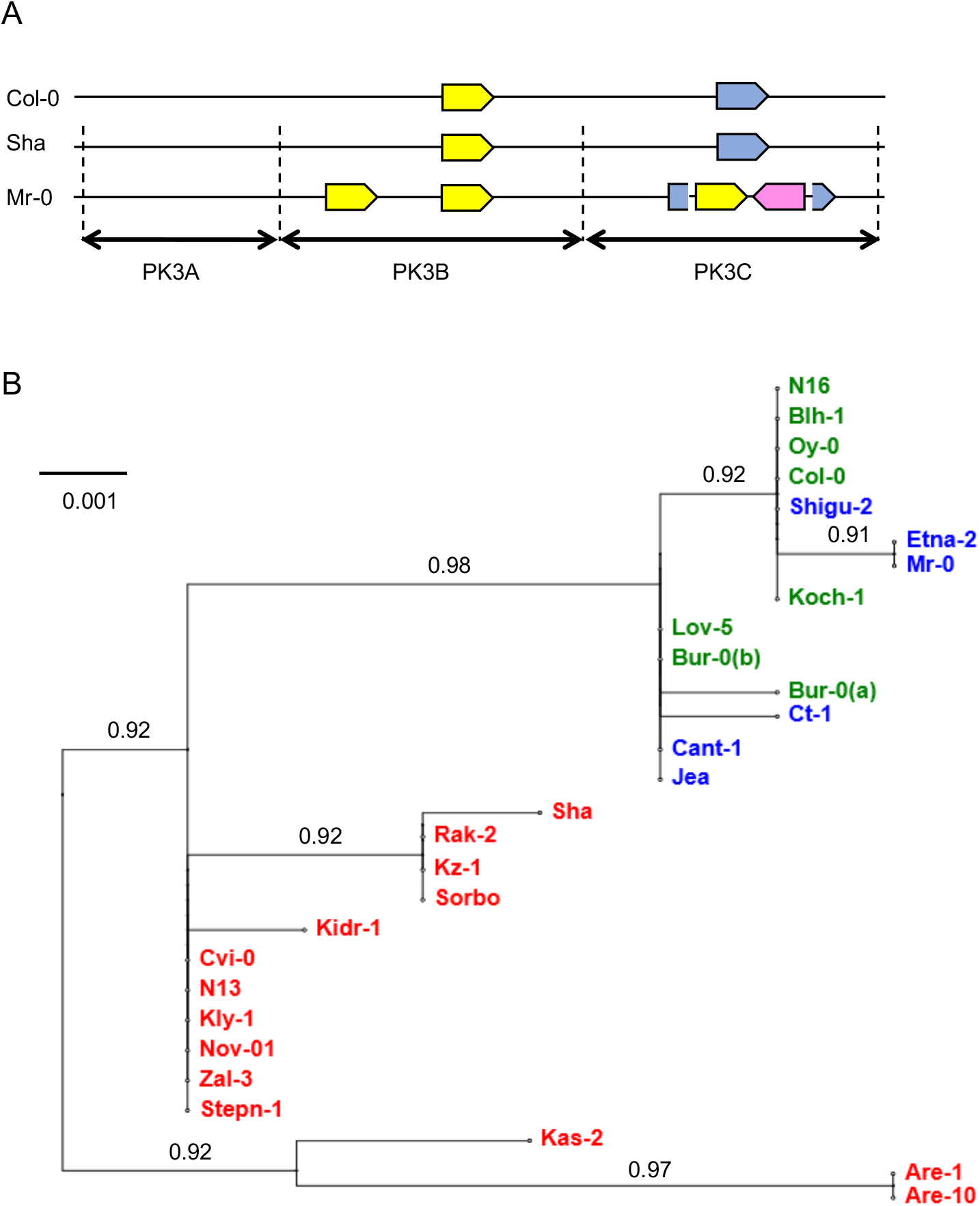
Variation in *APOK3* copy numbers and sequences. A: Schematic representation of *APOK3* homologous sequences (yellow arrows) in the PK3 locus in Col-0, Sha and Mr-0. The scheme is not to scale. Gene colours are as in Figure 7. The blue arrows represent *AT3G62610*, in which *APOK3-like* is inserted in Mr-0, together with *AT3G62540* (pink arrow). B: Variation in *APOK3* sequences : Unrooted phylogenic tree generated from the *APOK3* copies of 27 accessions of known status for the PK. DNA sequences were obtained after amplification with the primers AT3G62530F1 and AT3G62530R1 (Supplemental File 1) and the tree was generated using Phylogeny http://www.phylogeny.fr, (Dereeper et al. 2008) (Figure 10 Source Data 1). The names of killer, neutral and killed accessions are written in blue, green and red, respectively. The accessions with several copies of *APOK3* are displayed only once because their copies are identical, except Bur-0 where two sequences (a) differs from the third (b) by one SNP. Branch support values are displayed over the branches.

Our analysis of the natural variation at the PK3 locus revealed that all the killers analysed have multiple copies of *APOK3* (in yellow on Figure 7), with Mr-0 and Ct-1 having two copies, and Cant-1, Jea and Shigu-2 having three copies each. In addition, these five accessions have an *APOK3-like* copy inserted in *AT3G62610*. Bur-0, which has a neutral behavior, also possesses three copies of *APOK3*, but no *APOK3-like*, even if it has an insertion of *AT3G62540* in the intron of *AT3G62610*. The other neutral accessions and the killed ones have only one copy of *APOK3*, except Ita-0, in which no copy exists at the locus (Figure 7). No other copy of *APOK3* was found elsewhere in the Ita-0 genome, neither by searching in the genomic sequence nor by PCR amplification in Ita-0 genomic DNA. Similarly, we did not find any gene encoding a protein closer to APOK3 than to AT3G58180 neither in *A. lyrata* nor in other Brassicaceae sequences available in the databases. Altogether, these results suggest that *APOK3* is specific to *A. thaliana* and has evolved within the species.

### Antidote and non-antidote forms of APOK3 differ by three amino acids

In order to further explore the sequence variation of *APOK3* in relation with its antidote activity, we amplified and sequenced *APOK3* in all the accessions whose behavior for the PK3 had been determined (Table 1). The *APOK3* copies found in each killer accession whose genome was *de novo* sequenced were identical. In Etna-0, the only killer accession for which no genomic sequence was available, we obtained a unique Sanger sequence for *APOK3*, indicating that, whatever the number of copies it has, they are identical. Phylogenetic analysis clustered the sequences into three clades, one grouping all copies found in accessions possessing the antidote, *i*.*e*. killer and neutral. A second cluster groups most of copies from the killed accessions, with the exception of Are-1, Are-10 and Kas-2 which branch together as a third clade (Figure 10B). We identified 11 different *APOK3* haplotypes, five from non-killed accessions and six from killed accessions (Table 5). One polymorphism located 36 pb upstream of the ATG start codon and three non-synonymous SNPs in the coding sequence distinguished non-killed from killed alleles. Consequently, the APOK3 proteins with an antidote activity differ from those with no antidote activity by the three amino acid changes C85S, V101D and C105R (Table 5). Interestingly, the last two of these amino acids are located in the first HEAT-repeat domain of the protein, suggesting these changes could modify the protein interactions and could make it functional or non-functional as an antidote.

**Table 5:** APOK3 haplotype diversity among 27 accessions of known status for PK3

|  |  | SNP position<sup>(2)</sup> |  |  |  |  |  |  |  |  |  |  |  |  |  |  |  |  |  |  |  |
| --- | --- | --- | --- | --- | --- | --- | --- | --- | --- | --- | --- | --- | --- | --- | --- | --- | --- | --- | --- | --- | --- |
| Accessions <sup>(1)</sup> |  | -36 | 6 | 17 | 27 | 56 | 68 | 152 | 241 | 248 | 253 | 332 | 341 | 363 | 373 | 378 | 384 | 415 | 583 | 765 | 840 |
| antidotes | **Col-0, Shigu-2, Oy-0, Blh-1**, N16, Koch-1 | **c** | T | T | T | T | C | T | G | A | **A** | a | t | A | **A** | T | **C** | G | G | t | T |
|  | **Mr-0**, Etna-2 | . | . | . | . | . | . | . | T | . | . | . | . | . | . | . | . | . | . | . | . |
|  | **Ct-1** | . | . | . | . | . | . | . | . | . | . | . | . | T | . | . | . | . | . | g | . |
|  | **Bur-0 APOK3-1 & 2** | . | . | . | . | . | . | . | . | . | . | . | . | . | . | . | . | . | C | g | . |
|  | **Jea, Cant-1, Bur-0 APOK3-3**, Lov-5 | . | . | . | . | . | . | . | . | . | . | . | . | . | . | . | . | . | . | g | . |
| non-antidotes | **Cvi-0**, N13, Kly-1, Nov-01, Zal-3, Stepn-1 | **t** | . | . | . | . | . | . | . | . | **T** | . | . | . | **T** | . | **T** | . | . | g | . |
|  | Kidr-1 | **t** | . | . | . | . | . | . | . | . | **T** | . | c | . | **T** | . | **T** | . | . | g | . |
|  | Sorbo, Kz-1, Rak-2 | **t** | . | . | . | . | . | A | . | . | **T** | . | . | . | **T** | . | **T** | T | . | g | . |
|  | **Sha** | **t** | . | . | . | . | A | A | . | . | **T** | . | . | . | **T** | . | **T** | T | . | g | . |
|  | Kas-2 | **t** | . | . | . | C | . | . | . | C | **T** | . | . | . | **T** | G | **T** | . | . | g | G |
|  | **Are-10**, Are-1 | **g** | C | G | C | . | . | . | . | C | **T** | g | . | . | **T** | G | **T** | . | . | g | . |
| Amino acid change position |  |  |  | 6 |  | 19 | 23 | 51 | 81 | 83 | **85** |  |  | 98 | **101** | 103 | **105** | 115 | 141 |  |  |
| antidote |  |  |  | F |  | L | S | V | A | N | **S** |  |  | T | **D** | Y | **R** | G | V |  |  |
| non-antidote |  |  |  | C |  | S | Y | D | S | T | **C** |  |  | S | **V** | D | **C** | V | L |  |  |

<sup>(1)</sup> The names of the accessions whose entire PK3 sequences are known are in bold. The Col-0 sequence (TAIR 10) is used as reference. <sup>(2)</sup> Base 1 of the nucleotide sequence is the A of the start codon of the protein. The nucleotides in the exons are in upper case and those in the untranslated regions in lower case. Nucleotides identical to the reference sequence are represented as dots. Multiple sequence alignment was done with ClustalW (https://www.genome.jp/tools-bin/clustalw). SNPs and amino acids specific of antidote vs non-antidote forms are in bold.

## Discussion

As pointed out by Burga et al (2020), the discovery of poison-antidote systems in eukaryotes is most often fortuitous, especially because nearly all these systems are species specific. In *A. thaliana* too, we found PKs adventitiously when looking at a hybrid male sterility, of which PKs turned out to be unexpected components, in a cross between Sha and Mr-0 natural variants (Gobron et al. 2013; Simon et al. 2016). In this study, we focused on one of these PKs, responsible for a deficit in Sha homozygotes at the bottom of chromosome 3 in Mr-0/Sha hybrid progenies, due to the death of pollen grains carrying this allele. A TRD at the same locus was found in the progenies of 19 out of 44 hybrids involving either Sha or Mr-0 as one of the parents, indicating that killed or killer alleles are not rare in *A. thaliana* (Table 1). This is in agreement with the report by Seymour et al (2019) of frequent TRD in F2 populations from 80 *A. thaliana* founders, although these authors captured all types of segregation distorters, using a genome wide detection approach. Indeed, F2 distortion at one locus can result from allelic interactions either at the same locus or between different loci (Fishman and McIntosh 2019). We cannot formally exclude that some of the biases observed in our F2 families are partially caused by other allelic interactions than the PK3 studied here. However, our subsequent results on sequence variation in the antidote gene *APOK3*, which grouped antidote and non-antidote forms of the genes in distinct clades (Figure 10B), support our conclusion that we indeed detected the effect of PK3 in these crosses. In addition, in biased F2 populations, most killer:Hz ratios complied with the 1:1 ratio that is expected for a bias due to a gametophytic defect controlled by a single locus. The exceptions concerned the accessions Kas-2, Kly-1, N13 and Cvi-0 in crosses with Mr-0, for which F2 genotypes presented an excess in heterozygotes (>55%). These cases could be explained by the presence, at loci partially linked to the PK3, of other segregation distorters that affect the transmission of the Mr-0 allele.

Our cytological analyses showed that, in ShaL3^H^ plants, a defect of pollen development is visible as soon as the bicellular stage (Figure 4). The proportion of affected pollen then increases during development but reaches only approximately 35% of dead pollen in mature anthers instead of 50% expected if the killer effect was total. This incomplete penetrance of the killer effect is supported by the fact that plants homozygous for the killed allele are found in the progeny of heterozygotes. Indeed, if considering that all the 35% dead pollen carry a Sha allele, we theoretically expect 11,5% of Sha homozygotes in the progeny, which is very close to the 10% observed (Table 2). Incomplete penetrance was also observed in other reported gamete killers and meiotic drivers, for example in tomato (Rick 1966) or rice (Matsubara et al. 2011). In some cases, as in distorters from Drosophila (Larracuente and Presgraves 2012), rice (Koide et al. 2012) and yellow monkeyflower (Finseth et al. 2021), the incomplete penetrance of the phenotype was due to the presence of unlinked modifier loci that modulate the strength of the distorter. In this regard, it is interesting to note that the strength of the biases that we observed in the F2s vary between crosses, either with Sha or with Mr-0 (Table 1 Source Data 1 & 2). It is conceivable that this variation be due to the presence, in some hybrids, of unlinked epistatic loci involved in the sterility caused by PK3, even if in the case of Mr-0/Sha we observed no difference in the biases between plants with Sha and Mr-0 fixed nuclear backgrounds (Table 2). In contrast, changing the Mr-0 cytoplasmic background to the Sha one had a strong effect on the elimination of the Sha allele in pollen (Figure 9C), making it almost absolute (only one Sha homozygous plant among 512 progenies). Considering that APOK3 is a protein targeted to mitochondria, it is tempting to hypothesize that variations in the mitochondrial genome can modulate pollen sensitivity to the killer. In this case, the mitochondrial genome could be considered as a ‘modifier locus’ of the PK. Interestingly, the antidote of the rice *qHMS7* locus is also addressed to mitochondria (Yu et al. 2018) but the effect of the cytoplasmic background was not explored. To date, however, there is no indication that these poison-antidote systems from rice and Arabidopsis have functional similarities.

Our diversity survey revealed that some natural variants are neutral for the PK activity. Neutral alleles have been reported not only for gamete killers in rice (Koide et al. 2018; Yu et al. 2016; Wang et al. 2005; Liu et al. 2011) and tomato (Rick 1971), but also for spore killers in fungi such as Neurospora (Turner 2001) and for the meiotic driver *wtf* in fission yeast (Nuckolls et al. 2017). We took advantage of the neutral behavior of Col-0 to exploit available mutants affected in each of the protein coding genes present in the interval that contains the sensitivity factor of the PK. This allowed us to establish that the PK3 functions as a poison-antidote system and to identify the gene encoding the antidote, *APOK3*, as *AT3G62530*. Inactivation of *APOK3* in the neutral Col-0 allele turned it sensitive to a killer allele, as exemplified in hybrids with Mr-0 (Figure 8) and Ct-1 (Table 4), whereas hybrids with Col-0 or Sha had fully viable pollen and did not show any segregation distortion at the locus in their progenies (Table 4). These results demonstrate that the Col-0 allele of *APOK3* protects pollen from the effect of killer alleles in hybrids, which is the definition of an antidote. In addition, three residues in the protein sequence and one SNP in the 5’-UTR are strictly associated with the antidote forms of the gene (Table 5), suggesting their involvement in its protective activity.

APOK3 molecular function still remains to be elucidated. It is annotated belonging to the ARM-repeat superfamily, which mediate numerous cellular processes including signal transduction, cytoskeletal regulation, nuclear import, transcriptional regulation and ubiquitination (Samuel et al. 2006). The remarkable features of this protein are its interacting domains, its mitochondrial location and its chimeric structure. APOK3 has three HEAT repeat domains (Figure 9A), predicted to form super-helixes (Andrade and Bork 1995) and to mediate protein-protein interactions (Andrade et al. 2001). More specific investigations will be needed to determine if these domains are involved in interactions with other mitochondrial proteins. In this context, it is interesting to note that APOK3 was reported to associate with ISOCITRATE DEHYDROGENASE 1, a regulatory subunit of NAD+-dependent enzyme of the tricarboxylic acid cycle (TCA), and SUCCINATE DEHYDROGENASE 4, a subunit of the mitochondrial respiratory complex II, in a study exploring the protein-protein interaction network of the plant TCA cycle (Zhang et al. 2018). To date, the relevance to APOK3 function of its ability to bind zinc bivalent ions, reported by Tan et al. (Tan et al. 2010) is difficult to assess. However, these indications make excellent entry points to functional studies aiming at discovering how APOK3 fulfills its antidote activity. Indeed, this question has been elucidated in only very few systems, among which the *wtf* spore killer of fission yeast in which the antidote was shown to co-assemble with the corresponding poison and address the toxic aggregates to sequestering vacuoles (Nuckolls et al. 2020). Interestingly, these authors also found that genes involved in the mitochondrial functioning counteract the poison activity, suggesting a role, yet to be uncovered, of mitochondria in this antidote mechanism. From our results, it seems that APOK3 has no other biological role than being the antidote for PK3. In Col-0 the *apok3* KO mutation did not reveal any obvious phenotype alterations, at least in our laboratory standard conditions. Moreover, *APOK3* is missing in the Moroccan accession Ita-0. This supports that *APOK3* is not essential to *A. thaliana*. In addition to be absent in Ita-0, which belongs to an ancient lineage of *A. thaliana* (Arabidopsis Genome Consortium 2016; Durvasula et al. 2017), *APOK3* has no ortholog in the *A. thaliana* closest relative *A. lyrata* (Figure 7) and was not found in the available Brassicaceae sequences. This argues for this gene being specific to the *A. thaliana* species. It is thus likely that *APOK3* originated after *A. thaliana* divergence from its common ancestor with *A. lyrata*. The gene structure suggests that *APOK3* originated from a duplication of a member of the *AT3G43260* and *AT3G58180* gene family, followed by the recruitment of the *AT3G62460* mitochondrial targeting sequence. In *Oryza sativa* ssp *japonica*, a comparative genome analysis detected 28 new *O. sativa* specific genes on chromosome 3, and 14 of which were found to be chimerical (Zhang et al. 2013). However, as far as we are aware, their biological functions remain unknown. Because of the absence of *APOK3* in any related species, we could not conclusively infer whether non-antidote forms preceded antidote ones or rather derived from them. However, because *A. thaliana* Madeiran strains such as Are-1 and Are-10 have been described as archaic (Fulgione et al. 2018), our phylogenetic analysis of *APOK3* (Figure 10B) suggests that antidote forms evolved more recently. This would be an interesting question to address in order to understand the evolutionary trajectory of the gene in relation to its antidote function. Indeed, poison-antidote systems are often considered as selfish genetic elements, but their selfish nature is not always easy to properly establish, requiring thorough population and evolutionary genetic studies not amenable in all cases (Sweigart et al. 2019). In the present case, the question is still open, and deserves to be treated. It would involve the exploration of *APOK3* sequence diversity in a wider sampling of natural variants, in particular in polymorphic *A. thaliana* natural populations. The examination of the organization and gene content of 11 genotypes of different PK3 status in addition to Col-0, Sha and Mr-0 provide interesting information, though, revealing the strikingly complex and variable structure of the PK3 locus in particular in killer alleles. In eukaryotes, poison-antidote elements are generally found within regions of high divergence and structural variation (Burga et al. 2020). These are regions with low recombination (Larracuente and Presgraves 2012), which could favor the poison-antidote system. Indeed, the antidote must be inherited together with the killer. In our case, the structural differences between Sha and Mr-0 alleles easily explain the scarcity of recombinants found inside the PK3 interval during the fine-mapping of the locus. The use of these rare recombinants and of derived crosses allowed us to establish that the PK3 contains at least three genetic elements necessary for the PK activity (Figure 5), the antidote being flanked by two intervals each carrying elements necessary for the killer activity. Poison-antidote systems whose genetic factors have been identified so far in eukaryotes other than fungi most commonly involve two components (Burga et al. 2020). However, in plants, three components poison-antidote systems have been previously described in rice distorter loci. In rice S1 (Xie et al. 2019), three linked genes are necessary for toxicity; in rice S5 (Yang et al. 2012), two sporophytic factors are necessary for the killer activity but they are not carried by the same allele. From our mapping results, we cannot completely exclude that PK3A or PK3C also carry rescue activities not redundant with PK3B, nor that PK3B also carries killer activity not redundant with PK3A and PK3C. However, it is noticeable that this structure prevents the killer activity to be isolated from the antidote by a recombination event, since such an event would also separate the two mandatory killer elements.

The PK3 locus is particularly dynamic and prone to structural variations. Besides the Ita-0 allele that is structurally similar to that of *A. lyrata*, with orthologous genes in the same order and orientation (Figure 7), the other killed and the neutral alleles (excepted Bur-0) present the same organization of the locus (Figure 6B), even if some protein coding genes are missing in Are-10 and Cvi-0 (Figure 7). Among the neutral alleles, Bur-0 is an interesting exception: it resembles a killer both in structure (Figure 6B) and gene content (Figure 7), and we hypothesize it evolved from a killer allele that lost its killing capacity. Comparison of the Bur-0 sequence with the very similar allele of Jea will probably help identifying killer elements of PK3. The most complex alleles are found in killers (Figure 6B), which are highly variable, with different groups of genes duplicated in one or the other orientation and in different relative positions (Figure 7). Having more than one copy of *APOK3* is one remarkable common feature of the killer alleles. It is likely that having multiple copies is important for killer alleles not to be the victims of their own activity. Gene dosage has also been reported to be critical in a poison-antidote system in *C. elegans* (Mani and Fay 2009). In addition, the different copies of *APOK3* found in each killer are identical, but may be slightly different between accessions, indicating that recent duplications occurred independently in the lineages of the different variants we examined. These results suggest that these alleles have experienced a more intense structural evolution that neutral and killed ones, and raises the questions of the mechanisms and evolutive forces leading to these structures. The presence of several transposable elements at the locus, and their variation in occurrence between Sha, Mr-0 and Col-0 (Figure 6A) could be relevant to this question, since TE mobility is a main source of genetic variation in *A. thaliana* (Baduel et al. 2021). It has also been suggested that the proximity of transposons has facilitated duplication of the fission yeast *wtf* genes (Eickbush et al. 2019).

Even if gamete killers likely exist in all plant species, none had been investigated in *A. thaliana* until now, to our knowlege. The PK we dissected here has some general features of eukaryotic poison-antidote systems, including its species-specific nature and its presence within a hyper variable locus. The layout of the killer alleles is particular, with at least three mandatory elements and diverse duplications of sequence blocks that contain antidote genes trapped between killer elements. Continuing to exploit the natural variation of the species should help identifying the killer elements, and provide clues towards the underlying mechanisms responsible for the PK activity, the role of the mitochondria, and eventually the forces driving the evolution of the locus.

## Materials and Methods

### Plant material and growth conditions

*thaliana* natural accessions were provided by the Versailles Arabidopsis Stock Center (http://publiclines.versailles.inrae.fr/) and T-DNA lines were provided by the NASC (http://arabidopsis.info/).

In the analysis of the Sha/Mr bias at L3, all the genotypes used had the Mr-0 cytoplasm, unless specified. The ShaL3^H^ genotype was previously designated [Mr]ShaL3^H^ (Simon et al. 2016). It has the Mr-0 cytoplasmic genomes and a Sha nuclear background except at L3, *i*.*e*. between markers M1 and M15 (Figure 5 Source Data 3), where it is heterozygous. Here we constructed the MrL3^H^ genotype by backcrossing the (Mr-0 x Sha) F1 by Mr-0 three times. We selected a third backcross progeny heterozygous at L3, and at the loci L1 and L5 previously described in Simon et al (2016) and we chose in its self-descent a plant homozygous Mr-0 at L1 and L5 while heterozygous from the marker M0 (Figure 5 Source Data 3) to the bottom of chromosome 3, which we called MrL3^H^.

In order to save time, we constructed an early-flowering version of Mr-0 by inactivating *FRIGIDA* (*AT4G00650*) by CrispR-Cas9 (Doudna and Charpentier 2014). We cloned two guide-RNAs (275rev and 981forw, Supplemental File 1) in a pDe-Cas9-DsRed binary vector (Morineau et al. 2016) and introduced this construct by floral dipping (Clough and Bent 1998) in Mr-0 plants. We isolated several T1 transformants, some of them flowering earlier than Mr-0. After segregating out the T-DNA, we selected one T2 plant homozygous for a stop mutation in the *FRIGIDA* first exon. This early version of Mr-0, called Mr*fri*, flowers 39 days after sowing in our greenhouse conditions. Mr*fri* produces viable pollen, whereas its cross with Sha induces pollen lethality, as does Mr-0 (Figure 8 Supplement 1). As expected, it results in a segregation distortion against Sha alleles at L3 in the progeny of Mr*fri* x Sha plants.

Plants were grown in the greenhouse under long-day conditions (16h day, 8h night) with additional artificial light (105 µE/m2/sec) when necessary.

### Cytological analyses

DAPI staining of spread male meiotic chromosomes was performed according to (Ross et al. 1996). All stages were observed in 3 independent ShaL3^H^ plants and in parental controls. Observations were made using a Zeiss Axio Imager2 microscope and photographs were taken using an AxioCam MRm (Zeiss) camera. Propidium iodide and Alexander staining (Alexander 1969) of pollen were performed as described in Durand et al (2021).

### Fine-mapping and genotyping

DNA extractions were conducted on leaves from seedlings as described by Loudet et al (2002). Markers for the fine-mapping are described in Figure 5 Source Data 1. For CAPS markers M3 and M12 we used Cac8I (NEB) and Bsp1407I (ThermoFisher) restriction enzymes, respectively. Other SNPs were genotyped by sequencing. For the fine-mapping of PK3, we genotyped a total of 4,717 plants and identified 42 recombinants between markers M1 and M15. We selected 23 informative recombinants that were tested for segregation distortion at the locus by genotyping their self-descent progenies with appropriate markers (Figure 5 Source Data 1). All the other primers used in this work for mutant characterization, gene expression, PCR amplification and DNA sequencing are listed in Supplemental File 1.

### Purification of microspores

This step was carried out at the Imagerie-Gif facility (https://www.i2bc.paris-saclay.fr). Microspores were isolated from flower buds by chopping with a razor blade in 0.1M Mannitol. The crude suspension was filtered through a 50-µm nylon mesh (Sysmex-Partec) and collected in polypropylene tubes at 4°C. Microspores were sorted by flow cytometry using a MoFlo Astrios_EQ© cytometer (Beckman Coulter, Roissy, France) in PuraFlow sheath fluid (Beckman Coulter) at 25 psi (pounds per square inch), with a 100-micron nozzle. We performed sorting with ∼43 kHz drop drive frequency, plates voltage of 4000-4500 V and an amplitude of 30-50 V. Sorting was performed in purity mode. The 488-SSC (Side Scatter) parameter was set as threshold. Microspores gate was first set on FITC-A (526/50 band-pass for autofluorescence) versus SSC plot. Then, only low Forward Scatter (FSC) events were kept. The singlet gate was established using autofluorescence parameters. Accuracy of gating was determined post-sorting using microscopy with transmitted light and 40X dry. The flow cytometer-sorted microspores (300k) were collected in 1.5-ml tubes containing 300 µL Trireagent® and conserved at −80°C until subsequent RNA extraction.

### RNA extractions and RT-PCR

We mixed on ice 500 µl of microspore suspension in TriReagent with 300 µl glass beads (Sigma, G8772-100G glass beads acid washed) and two sterilized metals beads (3mm diameter). The tubes were shaked 2 × 2,5 min at a 1/30 frequency in a Retsch MM400 mixer mill and the solution without beads was recovered in a new RNase-free tube by making a small hole at the bottom of the tube and centrifuging 2 min at 3,900 rpm at 4°C. The beads were rinsed with 500 µl of TRIreagent and centrifuged again. The eluate was centrifuged 2min at 11,000 rpm, 4°C to remove cellular debris. RNA was extracted using the extraction RNA kit (Zymo research). Leaf RNA was prepared as above except that leaves were grinded in Trizol with one metal bead.

Reverse transcription was proceeded with 200 ng of purified RNA using the Maxima reverse transcriptase (Thermo Scientific) primed with oligodT in 50 µL. After heat inactivation of the RT, 2 µL of cDNA were used for each PCR.

### Sanger sequencing and annotation of Sha and Mr-0 PK3 locus

PCR and sequencing primers are listed in Supplemental File 1. Amplification products were sequenced by Beckman Coulter Genomics (http://www.beckmangenomics.com). Sequences were processed and aligned with Codon Code Aligner V5.0.2 (http://www.codoncode.com/aligner/). For Mr-0, because of the complexity of the locus, a fosmid library was built using the “Copycontrol fosmid library production kit” with pCC2FOS™ vector and T1 EPI300™ *E. coli* strain (Illumina technologies). The library titled 7.600 cfu/mL with an average size of inserts of 35 Kb. Purification of fosmid DNA was performed using the FosmidMax DNA purification kitTM (Illumina technologies).

The transfer of structural and functional annotation from the Col-0 reference genome to Sha and Mr-0 was done with the EGN-EP transfer pipeline (Sallet et al. 2019). These annotations were manually corrected, especially in Mr-0, to take into account copy number variations. TE annotation was performed with the TEannot tool (Quesneville et al. 2005), http://urgi.versailles.inra.fr/) and hits of more than 1 Kb were retained.

### *De novo* genome sequencing, assembly and annotation

High molecular weight DNA was extracted from 3 week-old plantlets using a protocol modified from Mayjonade et al (2016) which is described on Protocols.io (Russo et al. 2021). The following library preparation and sequencing were performed at the GeT-PlaGe core facility, INRAE Toulouse.

Nanopore sequencing: ONT libraries were prepared using the EXP-NBD103 and SQK-LSK109 kits according to the manufacturer’s instructions and using 4 µg of 40Kb sheared DNA (Megaruptor, Diagenode) as input. Pools of six samples were sequenced on one R9.4.1 flowcell. Between 0.014 and 0.020 pmol of library were loaded on each flowcell and sequenced on a PromethION instrument for 72 hours.

Illumina sequencing: Illumina libraries were prepared using the Illumina TruSeq Nano DNA HT Library Prep Kit according to the manufacter’s instructions. Libraries were then sequenced with 2×150bp paired-end reads on an Hiseq3000 instrument (Illumina).

Nanopore sequence datasets ranging from 4Gb to 14Gb with a minimum N50 of 22Kb were assembled with CANU (Koren et al. 2017) software release 1.9 (genomeSize=125M parameter). Mitochondrion and chloroplast genomes were assembled with CANU after selection of previously corrected CANU reads longer than 40Kb mapped with minimap2 (Li 2018) (–x asm5) on a databank built with Col-0 chloroplast and Col-0 and Landsberg mitochondrion genomes (NCBI accessions: NC_000932, NC_037304, JF729202). Spurious contigs were identified with minimap2 (-x asm5) and removed from the raw CANU assemblies. A spurious contig was defined as either a contig mapped on mitochondrion or chloroplast genomes or a contig mapped on a larger contig with a hit spanning at least 80% of its length. Nuclear contigs were scaffolded with AllMaps (Tang et al. 2015) using Col-0 genome as reference. Then, two rounds of consensus polishing using Illumina paired-end data were performed. For each round, the paired-end reads were first mapped with bwa (Li and Durbin 2009) (0.7.17-r1188 debian) with a minimal score of 50 (-T) then pilon software (Walker et al. 2014) (version 1.23) was used to generate the polished consensus sequences (--fix snps,indels --flank 20 –min-depth 20). Unanchored contigs shorter than 40kb were removed (considered as likely contaminants as very few Illumina reads were mapped on these short contigs).

The genomes were annotated using the EuGène software version 4.2a via the integrative pipeline egnep (Sallet et al. 2019) version 1.6 (http://eugene.toulouse.inra.fr/Downloads/egnep-Linux-x86_64.1.6.tar.gz). In addition to standard tools for repeat masking and lncRNA prediction included in egnep, the most recent and comprehensive annotation of *A. thaliana* available was used both as training dataset and as source of evidences. The peptide database of Araport (Krishnakumar et al. 2015) version 11 (201606) was used for the similarity searches performed with NCBI-BLASTX software version 2.2.31+ (hits spanning more than 80% of the protein length were retained). The corresponding cDNA database (201606) was used as transcriptional evidences. The cDNAs were mapped with gmap (Wu and Watanabe 2005) version 2017-09-05 (hits spanning more than 30% of the transcript length at a minimum identity percentage of 94 were retained).

In accessions with several copies of *APOK3*, occasional sequence errors subsist in some copies in the automatic assemblies. To correct these errors, *APOK3* was Sanger sequenced after gene amplification with the primers AT3G62530F1 and AT3G62530R1 (Supplemental File 1).

### Sequence availability

Whole genome sequences and PK3 locus sequences are available at the doi listed in Figure 6 Source Data 1. Start and end of the PK3 locus were determined by homology with the Col-0 reference sequence.

## Acknowledgments

We gratefully acknowledge Adeline Simon for her help in bioinformatics, Fabien Nogué for his advices in the CrispR Cas9 mutagenesis and Filipe Borges, Nicolas Valentin and Mickaël Bourge for the microspore FACS sorting. Sebastien Santini (CNRS/AMU IGS UMR7256) and the PACA Bioinfo platform (supported by IBISA) are acknowledged for the availability and management of the phylogeny.fr website. The present work has benefited from Imagerie-Gif core facility supported by l’Agence Nationale de la Recherche (ANR-11-EQPX-0029/Morphoscope, ANR-10-INBS-04/FranceBioImaging; ANR-11-IDEX-0003-02/ Saclay Plant Sciences) and from the support of IJPB Plant Observatory technological platforms. The IJPB benefits from the support of Saclay Plant Sciences-SPS (ANR-17-EUR-0007). This work was funded in part by INRAE Biology and Plant Breeding department (EVOLOX and POLLEN grants) and by the Région Midi-Pyrénées (CLIMARES project) and the LABEX TULIP (ANR-10-LABX-41).

## Competing interest statement

We declare no competing interests.

## Figure Supplements

Figure 8 Supplement 1: Mr*fri* has the same killer behaviour as Mr-0.

Figure 8 Supplement 2: Genetic and phenotypic characterization of the Col^mut530^ mutant.

Table 4 Supplement 1: Control of segregation at L3 in the progenies of F1 (Ct-1 x Col-0).

Figure 9 Supplement 1: Tree of protein sequences for deoxyhypusine hydroxylases from different taxa and their homologues in *A. thaliana*.

## Source Data Files

Table 1 Source Data 1: Segregations at L3 in progenies of crosses of Mr-0 with different natural accessions.

Table 2 Source Data 2: Segregations at L3 in progenies of crosses of different natural accessions with Sha.

Figure 4 Source Data 1: countings of abortive and normal grains in developing and mature pollen of plants with PK3 and parental controls.

Figure 5 Source Data 1: Segregation analyses in the self descent of 23 recombinants identified in PK3 fine mapping.

Figure 5 Source Data 2: Segregation at L3 in self progenies of hybrid plants derived from recombinants 52D7, 8F10BH2 and 23G9.

Figure 5 Source Data 3: Genetic markers.

Figure 6 Source Data 1: PK3 locus DNA sequence resources.

Figure 8 Source Data 1: annotated genes in the PK3B interval.

Figure 8 Source Data 2: Segregation analyses of Mrfri x Col^mut^ F2 families.

Figure 8 Source Data 3: Transmission of the Col^mut499^ and Col^mut530^ alleles from female and male sides.

Figure 8 Source Data 4: raw gels of RT-PCR experiments on PK3B genes.

Figure 9 Source Data 1: Segregation at L3 in reciprocal self progenies of hybrid plants derived from recombinants 52D7, 8F10BH2 and 18D1BC7.

Figure 10 Source Data 1: multifasta of APOK3 gene sequences from 27 accessions of knwon status at PK3.

Supplemental File 1: PCR primers.

**Figure 8 Supplement 1:**
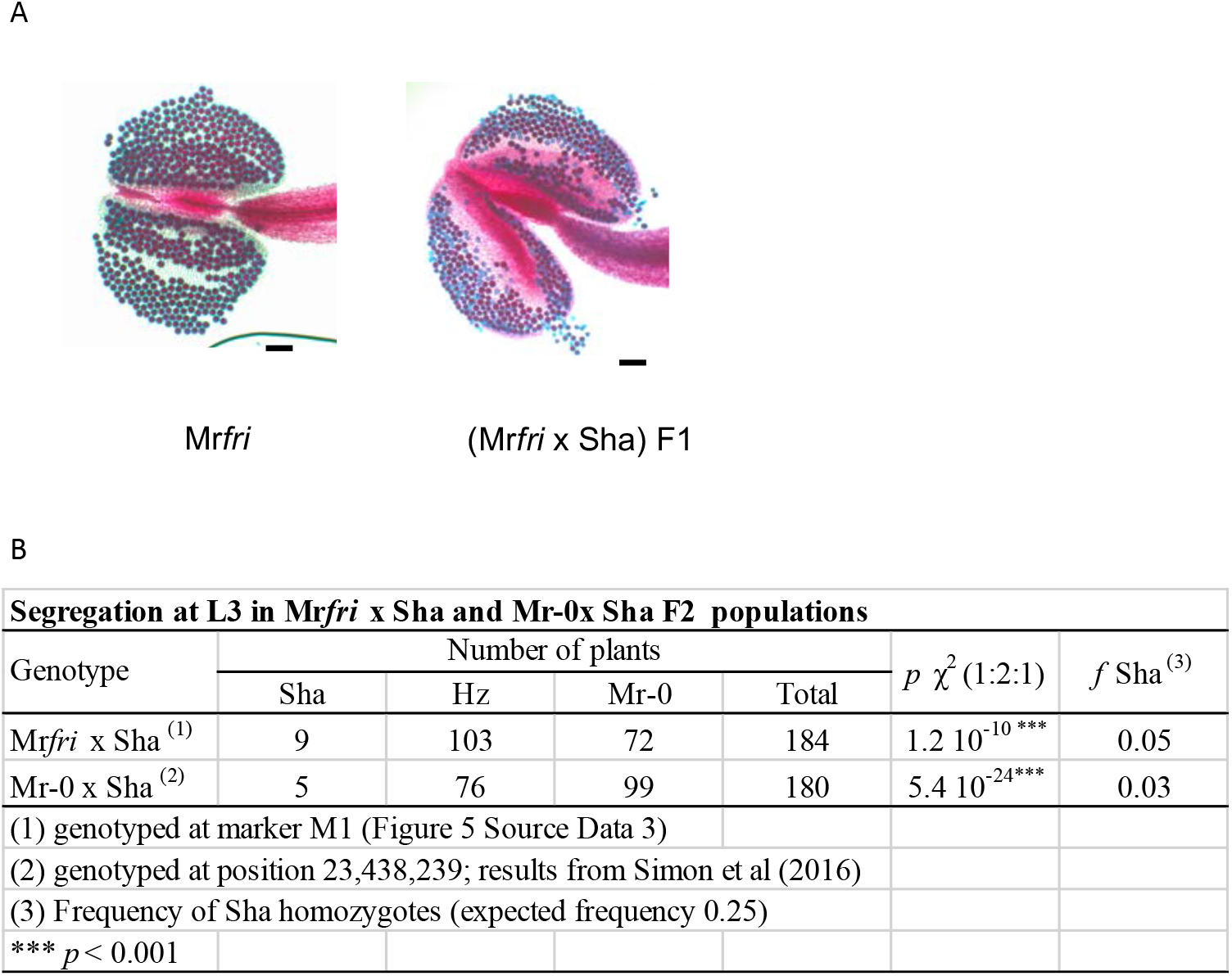
Mr*fri* has a killer behaviour at PK3, as Mr-0. (A) Pollen viability (Alexander stainings) in Mr*fri* plants and in the Mr*fri* x Sha F1. Viable pollen grains are stained in red, aborted pollen grains appear in blue. (B) Segregation at L3 in the selfed progeny of the Mr*fri* x Sha F1 showing the same transmission distortion as in the Mr-0 x Sha F1.

**Figure 8 supplement 2:**
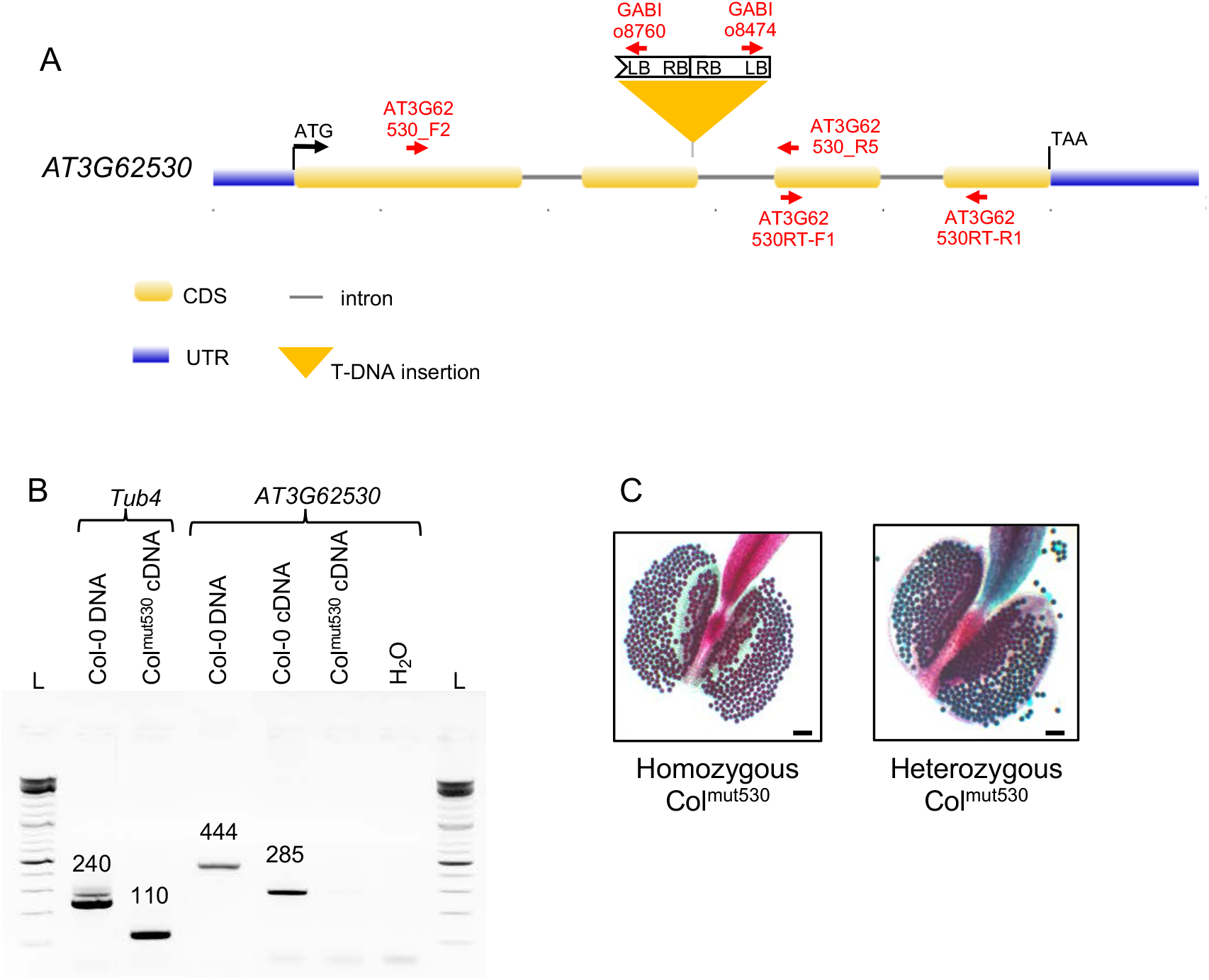
Genetic and phenotypic characterization of Col^mut530^. A: Representation of the position and schematic structure of GABI 745G01 T-DNA insertion in *AT3G62530*, with some primers (in red) used for mutant characterization and gene expression analysis. The T-DNA insertion is a reverse tandem with an incomplete LB border, located at the end of exon 2 (TAIR10 position 23132690). B: Expression analysis by RT-PCR of *AT3G62530* in wild type (Col-0) and homozygous mutant (Col^mut530^) plants. PCR primers used are AT3G62530RT-F1 & AT3G62530RT-R1 (Supplemental File 1). *Tub4* (*AT5G44340*) serves as a positive control. L: Thermo Scientific™ GeneRuler DNA ladder mix. The expected sizes of the amplification products are indicated in bp above the bands. *AT3G62530* is undetectable in cDNA from leaves of the homozygous mutant whereas it is detected in cDNA of Col-0 leaves. C: Representative images of anthers from homozygous and heterozygous Col^mut530^ plants after Alexander staining. Anthers present only viable pollen (in red), demonstrating the mutation by itself does not induce pollen abortion. Scale bars: 100µm.

**Figure 9 Supplement 1:**
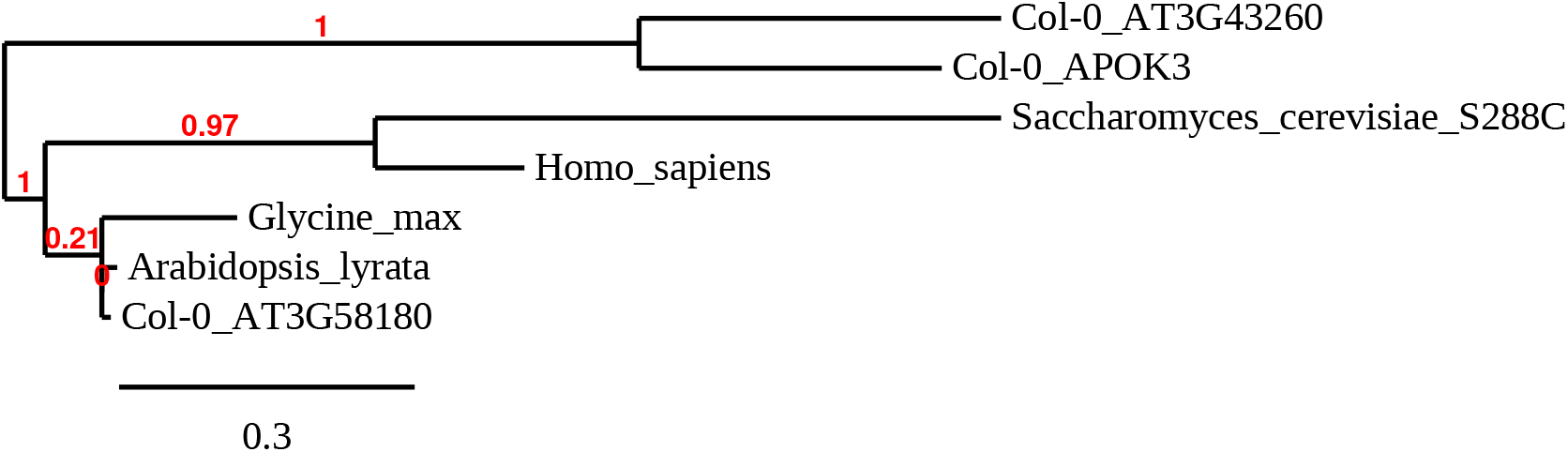
Tree of protein sequences for deoxyhypusine hydroxylases from different taxa and their homologues in *A. thaliana*. The hydroxyhypusine hydroxylase protein sequences from human (NP_001138637.1), baker’s yeast (*Saccharomyces cerevisiae*, NP_012604.1), soybean (*Glycine max*, XP_003521278.1), and *A. thaliana* sister species *A. lyrata* (XP_020880161.1) were aligned with those from Col-0 predicted products of genes *AT3G58180* (NP_567062.1), *AT3G43260* (NP_189912.1), and *AT3G62530* (APOK3; NP_567129.1), using Phylogeny.fr (http://www.phylogeny.fr/simple_phylogeny.cgi; (Dereeper et al. 2008) with defaults parameters. Branch support values are displayed in red over the branches.

## Notes

### Competing Interest Statement

The authors have declared no competing interest.

## References

Agorio A, Durand S, Fiume E, Brousse C, Gy I, Simon M, Anava S, Rechavi O, Loudet O, Camilleri C, Bouche N (2017) An Arabidopsis Natural Epiallele Maintained by a Feed-Forward Silencing Loop between Histone and DNA. PLoS Genet 13 (1):e1006551. doi:10.1371/journal.pgen.1006551

Agren JA, Clark AG (2018) Selfish genetic elements. PLoS Genet 14 (11):e1007700. doi:10.1371/journal.pgen.1007700

Alexander MP (1969) Differential staining of aborted and nonaborted pollen. Stain Technol 44 (3):117–122

Andrade MA, Bork P (1995) HEAT repeats in the Huntington’s disease protein. Nat Genet 11 (2):115–116. doi:10.1038/ng1095-115

Andrade MA, Petosa C, O’Donoghue SI, Muller CW, Bork P (2001) Comparison of ARM and HEAT protein repeats. J Mol Biol 309 (1):1–18. doi:10.1006/jmbi.2001.4624

Baduel P, Leduque B, Ignace A, Gy I, Gil J, Jr., Loudet O, Colot V, Quadrana L (2021) Genetic and environmental modulation of transposition shapes the evolutionary potential of Arabidopsis thaliana. Genome Biol 22 (1):138. doi:10.1186/s13059-021-02348-5

Bikard D, Patel D, Le Mette C, Giorgi V, Camilleri C, Bennett MJ, Loudet O (2009) Divergent evolution of duplicate genes leads to genetic incompatibilities within A. thaliana. Science 323 (5914):623–626. doi:10.1126/science.1165917

Bomblies K, Lempe J, Epple P, Warthmann N, Lanz C, Dangl JL, Weigel D (2007) Autoimmune response as a mechanism for a Dobzhansky-Muller-type incompatibility syndrome in plants. PLoS Biol 5 (9):e236. doi:07-PLBI-RA-1144 [pii] 10.1371/journal.pbio.0050236

Bravo Nunez MA, Nuckolls NL, Zanders SE (2018) Genetic Villains: Killer Meiotic Drivers. Trends Genet 34 (6):424–433. doi:10.1016/j.tig.2018.02.003

Burga A, Ben-David E, Kruglyak L (2020) Toxin-Antidote Elements Across the Tree of Life. Annu Rev Genet 54:387–415. doi:10.1146/annurev-genet-112618-043659

Clough SJ, Bent AF (1998) Floral dip: a simplified method for Agrobacterium-mediated transformation of Arabidopsis thaliana. Plant J 16 (6):735–743. doi:10.1046/j.1365-313x.1998.00343.x

Consortium” AG (2016) 1,135 Genomes Reveal the Global Pattern of Polymorphism in Arabidopsis thaliana. In: Cell, vol 166. vol 2, 2016/06/14 edn., pp 481–491. doi:10.1016/j.cell.2016.05.063

Doudna JA, Charpentier E (2014) Genome editing. The new frontier of genome engineering with CRISPR-Cas9. Science 346 (6213):1258096. doi:10.1126/science.1258096

Durand S, Bouche N, Perez Strand E, Loudet O, Camilleri C (2012) Rapid Establishment of Genetic Incompatibility through Natural Epigenetic Variation. Current biology : CB. doi:10.1016/j.cub.2011.12.054

Durand S, Ricou A, Simon M, Dehaene N, Budar F, Camilleri C (2021) A restorer-of-fertility-like pentatricopeptide repeat protein promotes cytoplasmic male sterility in Arabidopsis thaliana. Plant J 105 (1):124–135. doi:10.1111/tpj.15045

Durvasula A, Fulgione A, Gutaker RM, Alacakaptan SI, Flood PJ, Neto C, Tsuchimatsu T, Burbano HA, Pico FX, Alonso-Blanco C, Hancock AM (2017) African genomes illuminate the early history and transition to selfing in Arabidopsis thaliana. Proc Natl Acad Sci U S A 114 (20):5213–5218. doi:10.1073/pnas.1616736114

Eickbush MT, Young JM, Zanders SE (2019) Killer Meiotic Drive and Dynamic Evolution of the wtf Gene Family. Mol Biol Evol 36 (6):1201–1214. doi:10.1093/molbev/msz052

Finseth FR, Dong Y, Saunders A, Fishman L (2015) Duplication and Adaptive Evolution of a Key Centromeric Protein in Mimulus, a Genus with Female Meiotic Drive. Mol Biol Evol 32 (10):2694–2706. doi:10.1093/molbev/msv145

Finseth FR, Nelson TC, Fishman L (2021) Selfish chromosomal drive shapes recent centromeric histone evolution in monkeyflowers. PLoS Genet 17 (4):e1009418. doi:10.1371/journal.pgen.1009418

Fishman L, McIntosh M (2019) Standard Deviations: The Biological Bases of Transmission Ratio Distortion. Annu Rev Genet 53:347–372. doi:10.1146/annurev-genet-112618-043905

Fishman L, Sweigart AL (2018) When Two Rights Make a Wrong: The Evolutionary Genetics of Plant Hybrid Incompatibilities. Annu Rev Plant Biol 69:707–731. doi:10.1146/annurev-arplant-042817-040113

Fulgione A, Koornneef M, Roux F, Hermisson J, Hancock AM (2018) Madeiran Arabidopsis thaliana Reveals Ancient Long-Range Colonization and Clarifies Demography in Eurasia. Mol Biol Evol 35 (3):564–574. doi:10.1093/molbev/msx300

Gobron N, Waszczak C, Simon M, Hiard S, Boivin S, Charif D, Ducamp A, Wenes E, Budar F (2013) A cryptic cytoplasmic male sterility unveils a possible gynodioecious past for Arabidopsis thaliana. PLoS ONE 8 (4):e62450. doi:10.1371/journal.pone.0062450

Grognet P, Lalucque H, Malagnac F, Silar P (2014) Genes that bias Mendelian segregation. PLoS Genetics 10 (5):e1004387. doi:10.1371/journal.pgen.1004387

Heazlewood JL, Tonti-Filippini JS, Gout AM, Day DA, Whelan J, Millar AH (2004) Experimental analysis of the Arabidopsis mitochondrial proteome highlights signaling and regulatory components, provides assessment of targeting prediction programs, and indicates plant-specific mitochondrial proteins. Plant Cell 16 (1):241–256. doi:10.1105/tpc.016055

Houben A (2017) B Chromosomes - A Matter of Chromosome Drive. Front Plant Sci 8:210. doi:10.3389/fpls.2017.00210

Jiao WB, Patel V, Klasen J, Liu F, Pecinkova P, Ferrand M, Gy I, Camilleri C, Effgen S, Koornneef M, Pecinka A, Loudet O, Schneeberger K (2021) The Evolutionary Dynamics of Genetic Incompatibilities Introduced by Duplicated Genes in Arabidopsis thaliana. Mol Biol Evol 38 (4):1225–1240. doi:10.1093/molbev/msaa306

Klodmann J, Senkler M, Rode C, Braun HP (2011) Defining the protein complex proteome of plant mitochondria. Plant Physiol 157 (2):587–598. doi:10.1104/pp.111.182352

Koide Y, Ogino A, Yoshikawa T, Kitashima Y, Saito N, Kanaoka Y, Onishi K, Yoshitake Y, Tsukiyama T, Saito H, Teraishi M, Yamagata Y, Uemura A, Takagi H, Hayashi Y, Abe T, Fukuta Y, Okumoto Y, Kanazawa A (2018) Lineage-specific gene acquisition or loss is involved in interspecific hybrid sterility in rice. Proc Natl Acad Sci U S A 115 (9):E1955–E1962. doi:10.1073/pnas.1711656115

Koide Y, Shinya Y, Ikenaga M, Sawamura N, Matsubara K, Onishi K, Kanazawa A, Sano Y (2012) Complex genetic nature of sex-independent transmission ratio distortion in Asian rice species: the involvement of unlinked modifiers and sex-specific mechanisms. Heredity (Edinb) 108 (3):242–247. doi:10.1038/hdy.2011.64

Konig AC, Hartl M, Boersema PJ, Mann M, Finkemeier I (2014) The mitochondrial lysine acetylome of Arabidopsis. Mitochondrion 19 Pt B:252–260. doi:10.1016/j.mito.2014.03.004

Koren S, Walenz BP, Berlin K, Miller JR, Bergman NH, Phillippy AM (2017) Canu: scalable and accurate long-read assembly via adaptive k-mer weighting and repeat separation. Genome Res 27 (5):722–736. doi:10.1101/gr.215087.116

Krishnakumar V, Hanlon MR, Contrino S, Ferlanti ES, Karamycheva S, Kim M, Rosen BD, Cheng CY, Moreira W, Mock SA, Stubbs J, Sullivan JM, Krampis K, Miller JR, Micklem G, Vaughn M, Town CD (2015) Araport: the Arabidopsis information portal. Nucleic Acids Res 43 (Database issue):D1003–1009. doi:10.1093/nar/gku1200

Larracuente AM, Presgraves DC (2012) The Selfish Segregation Distorter Gene Complex of Drosophila melanogaster. Genetics 192 (1):33–53. doi:10.1534/genetics.112.141390

Li H (2018) Minimap2: pairwise alignment for nucleotide sequences. Bioinformatics 34 (18):3094–3100. doi:10.1093/bioinformatics/bty191

Li H, Durbin R (2009) Fast and accurate short read alignment with Burrows-Wheeler transform. Bioinformatics 25 (14):1754–1760. doi:10.1093/bioinformatics/btp324

Lindholm AK, Dyer KA, Firman RC, Fishman L, Forstmeier W, Holman L, Johannesson H, Knief U, Kokko H, Larracuente AM, Manser A, Montchamp-Moreau C, Petrosyan VG, Pomiankowski A, Presgraves DC, Safronova LD, Sutter A, Unckless RL, Verspoor RL, Wedell N, Wilkinson GS, Price TAR (2016) The Ecology and Evolutionary Dynamics of Meiotic Drive. Trends Ecol Evol 31 (4):315–326. doi:10.1016/j.tree.2016.02.001

Liu B, Li JQ, Liu XD, Shahid MQ, Shi LG, Lu YG (2011) Identification of neutral genes at pollen sterility loci Sd and Se of cultivated rice (Oryza sativa) with wild rice (O. rufipogon) origin. Genet Mol Res 10 (4):3435–3445. doi:10.4238/2011.October.31.10

Long Y, Zhao L, Niu B, Su J, Wu H, Chen Y, Zhang Q, Guo J, Zhuang C, Mei M, Xia J, Wang L, Liu YG (2008) Hybrid male sterility in rice controlled by interaction between divergent alleles of two adjacent genes. Proc Natl Acad Sci U S A 105 (48):18871–18876. doi:10.1073/pnas.0810108105

Loudet O, Chaillou S, Camilleri C, Bouchez D, Daniel-Vedele F (2002) Bay-0 x Shahdara recombinant inbred line population: a powerful tool for the genetic dissection of complex traits in Arabidopsis. Theor Appl Genet 104 (6-7):1173–1184. doi:10.1007/s00122-001-0825-9

Mani K, Fay DS (2009) A mechanistic basis for the coordinated regulation of pharyngeal morphogenesis in Caenorhabditis elegans by LIN-35/Rb and UBC-18-ARI-1. PLoS Genet 5 (6):e1000510. doi:10.1371/journal.pgen.1000510

Matsubara K, Ebana K, Mizubayashi T, Itoh S, Ando T, Nonoue Y, Ono N, Shibaya T, Ogiso E, Hori K, Fukuoka S, Yano M (2011) Relationship between transmission ratio distortion and genetic divergence in intraspecific rice crosses. Mol Genet Genomics 286 (5-6):307–319. doi:10.1007/s00438-011-0648-6

Mayjonade B, Gouzy J, Donnadieu C, Pouilly N, Marande W, Callot C, Langlade N, Munos S (2016) Extraction of high-molecular-weight genomic DNA for long-read sequencing of single molecules. Biotechniques 61 (4):203–205. doi:10.2144/000114460

Morineau C, Gissot L, Bellec Y, Hematy K, Tellier F, Renne C, Haslam R, Beaudoin F, Napier J, Faure JD (2016) Dual Fatty Acid Elongase Complex Interactions in Arabidopsis. PLoS One 11 (9):e0160631. doi:10.1371/journal.pone.0160631

Nuckolls NL, Bravo Nunez MA, Eickbush MT, Young JM, Lange JJ, Yu JS, Smith GR, Jaspersen SL, Malik HS, Zanders SE (2017) wtf genes are prolific dual poison-antidote meiotic drivers. Elife 6. doi:10.7554/eLife.26033

Nuckolls NL, Mok AC, Lange JJ, Yi K, Kandola TS, Hunn AM, McCroskey S, Snyder JL, Bravo Nunez MA, McClain M, McKinney SA, Wood C, Halfmann R, Zanders SE (2020) The wtf4 meiotic driver utilizes controlled protein aggregation to generate selective cell death. Elife 9. doi:10.7554/eLife.55694

Östergren G (1945) Parasitic nature of extra fragment chromosomes. Bot Not 2:157–163

Ouyang Y, Zhang Q (2013) Understanding reproductive isolation based on the rice model. Annu Rev Plant Biol 64:111–135. doi:10.1146/annurev-arplant-050312-120205

Ouyang Y, Zhang Q (2018) The molecular and evolutionary basis of reproductive isolation in plants. J Genet Genomics 45 (11):613–620. doi:10.1016/j.jgg.2018.10.004

Presgraves DC (2010) The molecular evolutionary basis of species formation. Nature reviews Genetics 11 (3):175–180. doi:10.1038/nrg2718

Quesneville H, Bergman CM, Andrieu O, Autard D, Nouaud D, Ashburner M, Anxolabehere D (2005) Combined evidence annotation of transposable elements in genome sequences. PLoS Comput Biol 1 (2):166–175. doi:10.1371/journal.pcbi.0010022

Rick CM (1966) Abortion of male and female gametes in the tomato determined by allelic interaction. Genetics 53 (1):85–96

Rick CM (1971) The tomato ge locus: linkage relations and geographic distribution of alleles. Genetics 67 (1):75–85

Ross KJ, Fransz P, Jones GH (1996) A light microscopic atlas of meiosis in Arabidopsis thaliana. Chromosome Res 4 (7):507–516. doi:10.1007/BF02261778

Russo A, Potente G, Mayjonade B (2021) HMW DNA extraction from diverse plants species for PacBio and Nanopore sequencing. protocolsio doi:dx.doi.org/10.17504/protocols.io.5t7g6rn

Sallet E, Gouzy J, Schiex T (2019) EuGene: An Automated Integrative Gene Finder for Eukaryotes and Prokaryotes. Methods Mol Biol 1962:97–120. doi:10.1007/978-1-4939-9173-0_6

Salome PA, Bomblies K, Fitz J, Laitinen RA, Warthmann N, Yant L, Weigel D (2012) The recombination landscape in Arabidopsis thaliana F2 populations. Heredity (Edinb) 108 (4):447–455. doi:10.1038/hdy.2011.95

Samuel MA, Salt JN, Shiu SH, Goring DR (2006) Multifunctional arm repeat domains in plants. Int Rev Cytol 253:1–26. doi:10.1016/S0074-7696(06)53001-3

Senkler J, Senkler M, Eubel H, Hildebrandt T, Lengwenus C, Schertl P, Schwarzlander M, Wagner S, Wittig I, Braun HP (2017) The mitochondrial complexome of Arabidopsis thaliana. Plant J 89 (6):1079–1092. doi:10.1111/tpj.13448

Seymour DK, Chae E, Arioz BI, Koenig D, Weigel D (2019) Transmission ratio distortion is frequent in Arabidopsis thaliana controlled crosses. Heredity (Edinb) 122 (3):294–304. doi:10.1038/s41437-018-0107-9

Simon M, Durand S, Pluta N, Gobron N, Botran L, Ricou A, Camilleri C, Budar F (2016) Genomic Conflicts that Cause Pollen Mortality and Raise Reproductive Barriers in Arabidopsis thaliana. Genetics 203 (3):1353–1367. doi:10.1534/genetics.115.183707

Simon M, Simon A, Martins F, Botran L, Tisne S, Granier F, Loudet O, Camilleri C (2012) DNA fingerprinting and new tools for fine-scale discrimination of Arabidopsis thaliana accessions. The Plant journal : for cell and molecular biology 69 (6):1094–1101. doi:10.1111/j.1365-313X.2011.04852.x

Sweigart AL, Brandvain Y, Fishman L (2019) Making a Murderer: The Evolutionary Framing of Hybrid Gamete-Killers. Trends Genet. doi:10.1016/j.tig.2019.01.004

Tan YF, O’Toole N, Taylor NL, Millar AH (2010) Divalent metal ions in plant mitochondria and their role in interactions with proteins and oxidative stress-induced damage to respiratory function. Plant Physiol 152 (2):747–761. doi:10.1104/pp.109.147942

Tang H, Zhang X, Miao C, Zhang J, Ming R, Schnable JC, Schnable PS, Lyons E, Lu J (2015) ALLMAPS: robust scaffold ordering based on multiple maps. Genome Biol 16:3. doi:10.1186/s13059-014-0573-1

Taylor NL, Heazlewood JL, Millar AH (2011) The Arabidopsis thaliana 2-D gel mitochondrial proteome: Refining the value of reference maps for assessing protein abundance, contaminants and post-translational modifications. Proteomics 11 (9):1720–1733. doi:10.1002/pmic.201000620

Turner BC (2001) Geographic distribution of neurospora spore killer strains and strains resistant to killing. Fungal Genet Biol 32 (2):93–104. doi:10.1006/fgbi.2001.1253

Vaid N, Laitinen RAE (2019) Diverse paths to hybrid incompatibility in Arabidopsis. Plant J 97 (1):199–213. doi:10.1111/tpj.14061

Walker BJ, Abeel T, Shea T, Priest M, Abouelliel A, Sakthikumar S, Cuomo CA, Zeng Q, Wortman J, Young SK, Earl AM (2014) Pilon: an integrated tool for comprehensive microbial variant detection and genome assembly improvement. PLoS One 9 (11):e112963. doi:10.1371/journal.pone.0112963

Wang GW, He YQ, Xu CG, Zhang Q (2005) Identification and confirmation of three neutral alleles conferring wide compatibility in inter-subspecific hybrids of rice (Oryza sativa L.) using near-isogenic lines. Theor Appl Genet 111 (4):702–710. doi:10.1007/s00122-005-2055-z

Wu TD, Watanabe CK (2005) GMAP: a genomic mapping and alignment program for mRNA and EST sequences. Bioinformatics 21 (9):1859–1875. doi:10.1093/bioinformatics/bti310

Xie Y, Tang J, Xie X, Li X, Huang J, Fei Y, Han J, Chen S, Tang H, Zhao X, Tao D, Xu P, Liu YG, Chen L (2019) An asymmetric allelic interaction drives allele transmission bias in interspecific rice hybrids. Nat Commun 10 (1):2501. doi:10.1038/s41467-019-10488-3

Yang J, Zhao X, Cheng K, Du H, Ouyang Y, Chen J, Qiu S, Huang J, Jiang Y, Jiang L, Ding J, Wang J, Xu C, Li X, Zhang Q (2012) A killer-protector system regulates both hybrid sterility and segregation distortion in rice. Science 337 (6100):1336–1340. doi:10.1126/science.1223702

Yu X, Zhao Z, Zheng X, Zhou J, Kong W, Wang P, Bai W, Zheng H, Zhang H, Li J, Liu J, Wang Q, Zhang L, Liu K, Yu Y, Guo X, Wang J, Lin Q, Wu F, Ren Y, Zhu S, Zhang X, Cheng Z, Lei C, Liu S, Liu X, Tian Y, Jiang L, Ge S, Wu C, Tao D, Wang H, Wan J (2018) A selfish genetic element confers non-Mendelian inheritance in rice. Science 360 (6393):1130–1132. doi:10.1126/science.aar4279

Yu Y, Zhao Z, Shi Y, Tian H, Liu L, Bian X, Xu Y, Zheng X, Gan L, Shen Y, Wang C, Yu X, Wang C, Zhang X, Guo X, Wang J, Ikehashi H, Jiang L, Wan J (2016) Hybrid Sterility in Rice (Oryza sativa L.) Involves the Tetratricopeptide Repeat Domain Containing Protein. Genetics 203 (3):1439–1451. doi:10.1534/genetics.115.183848

Zhang C, Wang D, Wang J, Sun Q, Tian L, Tang X, Yuan Z, He H, Yu S (2020) Genetic Dissection and Validation of Chromosomal Regions for Transmission Ratio Distortion in Intersubspecific Crosses of Rice. Front Plant Sci 11:563548. doi:10.3389/fpls.2020.563548

Zhang C, Wang J, Marowsky NC, Long M, Wing RA, Fan C (2013) High occurrence of functional new chimeric genes in survey of rice chromosome 3 short arm genome sequences. Genome Biol Evol 5 (5):1038–1048. doi:10.1093/gbe/evt071

Zhang Y, Swart C, Alseekh S, Scossa F, Jiang L, Obata T, Graf A, Fernie AR (2018) The Extra-Pathway Interactome of the TCA Cycle: Expected and Unexpected Metabolic Interactions. Plant Physiol 177 (3):966–979. doi:10.1104/pp.17.01687

